# Vision-dependent specification of cell types and function in the developing cortex

**DOI:** 10.1101/2021.08.10.455824

**Authors:** Sarah Cheng, Salwan Butrus, Liming Tan, Vincent Xu, Srikant Sagireddy, Joshua T. Trachtenberg, Karthik Shekhar, S. Lawrence Zipursky

## Abstract

The role of postnatal experience in sculpting cortical circuitry, while long appreciated, is poorly understood at the level of cell types. We explore this in the mouse primary visual cortex (V1) using single-nucleus RNA-sequencing, visual deprivation, genetics, and functional imaging. We find that vision selectively drives the specification of glutamatergic cell types in upper layers (L) (L2/3/4), while deeper-layer glutamatergic, GABAergic, and non-neuronal cell types are established prior to eye opening. L2/3 cell types form an experience-dependent spatial continuum defined by the graded expression of ∼200 genes, including regulators of cell adhesion and synapse formation. Vision-dependent regulation of one of these genes, encoding the inhibitory synaptic cell adhesion molecule IGSF9b, is required for the normal development of binocular responses in L2/3. In summary, vision preferentially regulates the development of upper-layer glutamatergic cell types through the regulation of cell type-specific gene expression programs.

## INTRODUCTION

The establishment of neural circuitry in the mammalian cortex relies on the interaction of the developing postnatal animal with its environment. Cortical circuits comprise diverse cell types and a network of complex patterns of synaptic connections between them ^1,2^. The formation of this circuitry relies on genetically hard-wired mechanisms mediated by cell recognition molecules and sensory-independent neural activity ^3–8^. During postnatal development, experience-dependent processes are required for the maturation of this circuitry ^5,9–12^. These periods of developmental plasticity, known as “critical periods”, are observed in sensory cortical areas and regulate more complex multimodal processes such as language development and cognition ^13^.

The influence of experience on cortical circuitry in the primary visual cortex (V1) is accessible to molecular, genetic, and functional analysis in the mouse and thus is particularly well suited for mechanistic studies ^9^. Neural circuitry is patterned by vision ^14,15^ and this process can be studied through longitudinal calcium imaging of neurons in V1 of awake behaving animals ^15,16^. Mice open their eyes around postnatal day (P)14. Binocular circuitry is sensitive to vision immediately after eye opening, but its peak period of sensitivity, demonstrated and defined by the effects of monocular deprivation on cortical ocular dominance, begins about a week after eye opening (∼P21) and continues through ∼P35 ^14,17^. Visual experience during this period is necessary for the development and maintenance of the neural circuitry underlying binocular vision ^5,14–19^.

Recent advances in single-cell transcriptomics have uncovered a vast diversity of neuronal cell types in the adult mouse V1 ^1,20,21^. Previous investigations of vision-dependent changes in gene expression during the critical period have relied on comparing bulk transcriptomic profiles of V1 between normally reared and visually deprived animals ^22–24^, or within normally reared animals at different points during the critical period ^25^. Consequently, these studies did not investigate vision-dependent gene expression at the level of the diverse cell types in V1. This resolution is crucial to enable mechanistic understanding of experience-dependent regulation of neural circuitry at the molecular, cellular, and functional levels.

Here, we study the role of vision in the development of V1 cell types and their circuitry in mice by combining single-cell transcriptomics, statistical inference, sensory perturbations, genetics, and *in vivo* functional imaging. We used single-nucleus RNA-seq to construct a developmental atlas of cell types in mouse V1 during early postnatal life in normal and dark-reared animals. Through computational analysis of cell type maturation, we discover that glutamatergic (excitatory) cell type identities in superficial layers (L2/3/4) are specified after eye opening, whereas the identities of deeper-layer glutamatergic neurons, GABAergic neurons and non-neuronal cell types are established before eye opening. We report that L2/3 glutamatergic cell types are organized as sublayers in V1 and form a transcriptomic continuum through the graded expression of ∼200 genes. This continuum relies on vision for its establishment and maintenance during the critical period. Through genetics and live imaging of visually evoked activity in alert mice, we show that vision acts through one of these genes, *Igsf9b*, encoding a homophilic cell adhesion protein that regulates inhibitory synapse formation. Deletion of this gene impairs the maturation of neural circuitry during the critical period in a spatially graded fashion. Thus, one way that visual experience regulates the specification of cell types and sculpts function in V1 is by regulating intercellular adhesion and inhibitory synapses.

## RESULTS

### Transcriptional profiling of mouse V1 development using single-nucleus RNA-seq

To survey the transcriptomic diversity and maturation of cells in V1, we used droplet-based single-nucleus (sn) RNA-seq to profile this region during postnatal development in normally reared mice (Figures 1A and S1D). We collected samples from six postnatal time points: P8, P14, P17, P21, P28 and P38 (Figure 1B). The first three time points are prior to the classical critical period for ocular dominance plasticity, with synaptogenesis occurring between P8 and eye-opening (P14) ^26,27^ (Figures S1A-C). The final three timepoints cover the critical period of ocular dominance plasticity ^14,15,18^ including its start (P21), peak (P28), and closure (P38).

**Figure 1.**
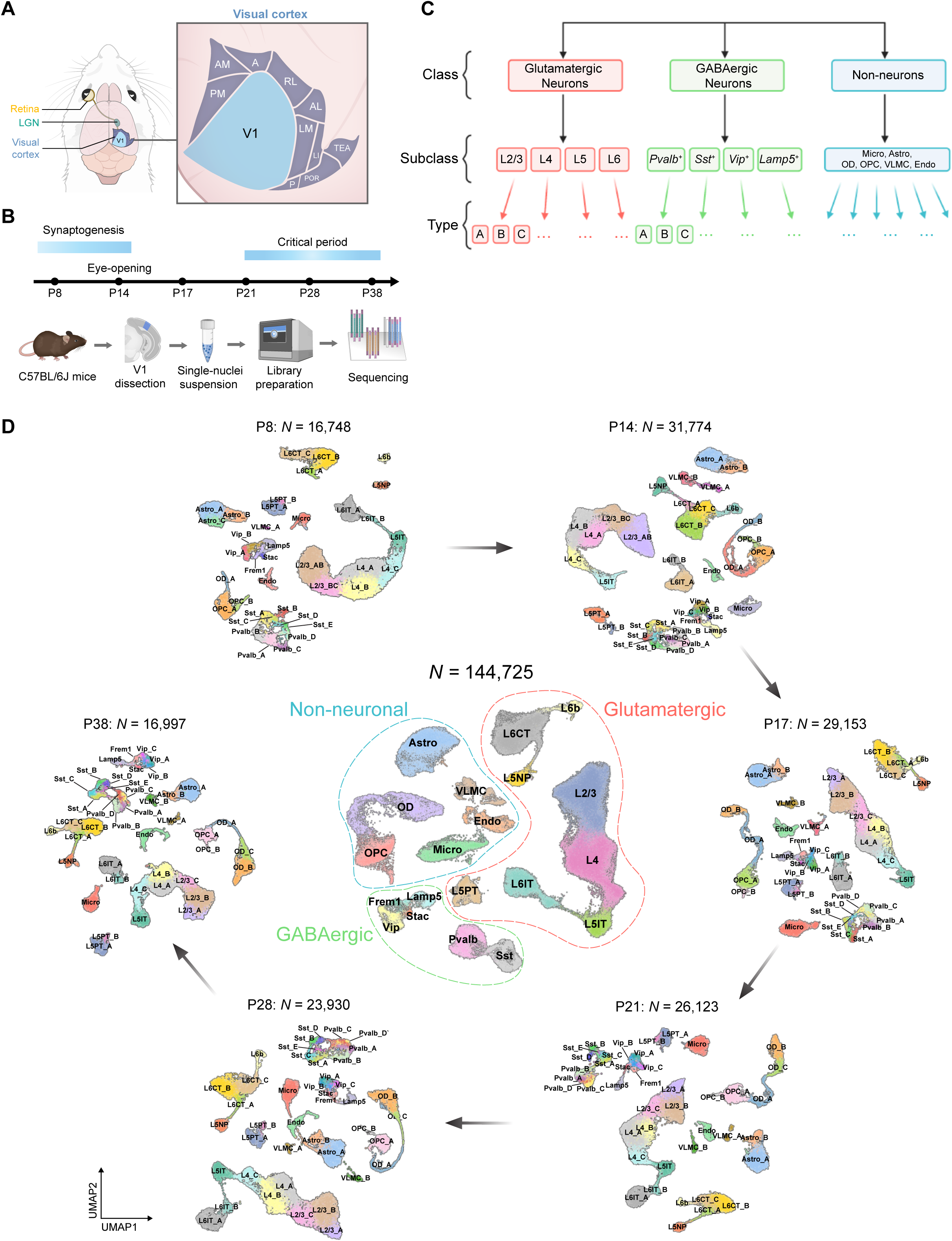
snRNA-seq profiling of V1 during postnatal development. A. Schematic of the mouse visual system. Visual information predominantly passes from the retina to the contralateral visual cortex through the lateral geniculate nucleus (LGN). Some visual information from the ipsilateral eye also passes through the LGN to the ipsilateral cortex (not shown). Inset shows a magnified view of the primary visual cortex (V1) and surrounding higher visual areas. Abbreviations: A, anterior; AL, anterolateral; AM, anteromedial; LI, laterointermediate; LM, lateromedial; P, posterior; PM, posteromedial; POR, postrhinal; RL, rostrolateral; TEA, temporal anterior areas. B. Experimental workflow of snRNA-seq profiling of V1 at six postnatal ages (see **Methods** for details). C. Cellular taxonomy of V1 showing the hierarchical organization of transcriptomic classes, subclasses, and types. D. Uniform Manifold Approximation and Projection (UMAP) visualization of V1 transcriptomic diversity during postnatal development ^56^. Dots correspond to cells and distances between them reflect degrees of transcriptomic similarity. Central panel shows cells from all six ages colored by subclass identity (**Table S1**). The six peripheral panels show cells from different ages, colored by type identity determined via clustering. Data from each age and class were analyzed separately and then merged together for visualization purposes (**Methods**).

Data from each timepoint consisted of four single-nuclei library replicates, each derived from cells collected from multiple mice (**Methods**). The resulting gene expression matrices were filtered to remove low-quality cells and doublets ^28^, as well as cells with a high proportion of mitochondrial transcripts (>1%). In total, we obtained 144,725 high-quality nuclear transcriptomes across the six time points (Figures S1D-H).

### A postnatal developmental atlas of V1 cell classes, subclasses, and types

We used dimensionality reduction and clustering to derive a developmental taxonomy consisting of cell classes, subclasses, and types ^29–31^ at each of the six time points (Figures 1C, D and S1D**; Methods**). Cell classes consisted of glutamatergic neurons (n=92,856; 3176 genes/cell detected on average), GABAergic neurons (n=13,374; 2966 genes/cell), and non-neuronal cells (n=38,495; 1549 genes/cell) identified by canonical markers (Figure S1I and **Table S1**) ^1,20,21^. The relative proportions of the three cell classes were consistent across biological replicates (data not shown).

Glutamatergic cells separated into eight subclasses within the four cortical layers - L2/3, L4, L5IT, L5NP, L5PT, L6CT, L6IT, and L6b (Figures 2A, B). We also identified six GABAergic subclasses, which included the four well-known subclasses defined by the selective expression of *Pvalb*, *Sst*, *Vip*, and *Lamp5* ^29^ and two smaller subclasses that selectively expressed the genes *Stac* and *Frem1*. Non-neuronal cells included oligodendrocytes, oligodendrocyte precursor cells, astrocytes, vascular and leptomeningeal cells, endothelial cells, and microglia (Figure 1D). Similar results were obtained using an alternative computational pipeline ^32^ (Figure S1K). We found a tight correspondence between the transcriptome-wide gene signatures that defined developing subclasses in our dataset and the subclasses identified in a recent survey of the adult mouse cortex ^20^ (Figure S1J).

**Figure 2.**
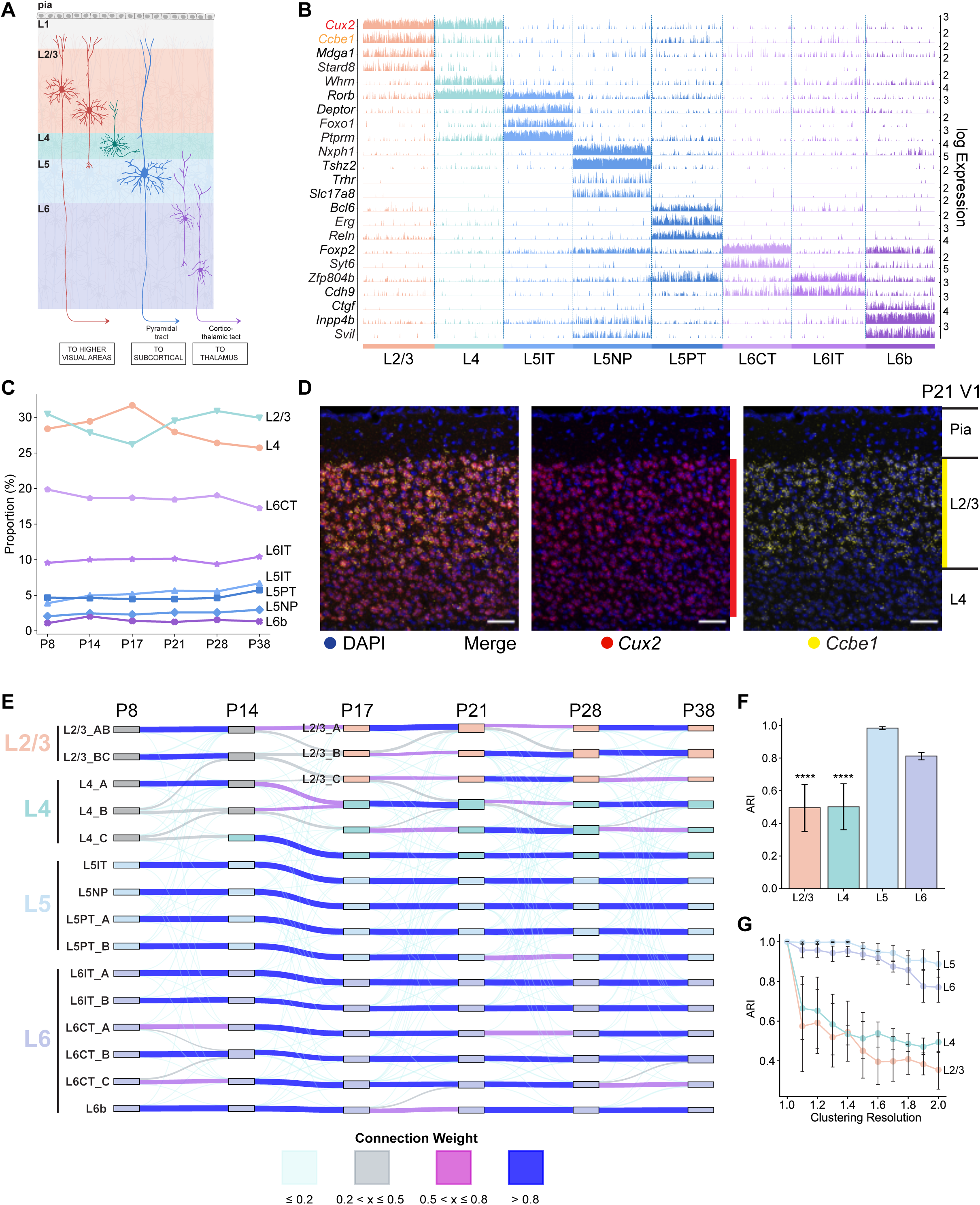
Transcriptomic diversity of V1 glutamatergic neurons during postnatal development. A. Schematic of glutamatergic neurons in V1 arranged in layers L1-L6. B. Tracks plot showing subclass-specific markers (rows) in glutamatergic neurons (columns), grouped by subclass (annotation bar, bottom). 1000 randomly selected cells from each subclass were used for plotting. For each gene, the scale on the y-axis (right) corresponds to normalized, log-transformed transcript counts detected in each cell. *Ccbe1*, a L2/3 marker, and *Cux2*, a L2/3/4 marker, are highlighted. C. Line plots showing that the proportions of glutamatergic subclasses are stable with age despite significant variation in the number of cells profiled (**Table S2**). D. *Ccbe1* is a selective marker for L2/3 glutamatergic neurons. By contrast, *Cux2*, which is expressed in L2/3 and L4 glutamatergic neurons, is also expressed in inhibitory neurons and non-neuronal cells (also see Figure S2B). Panels show a coronal section through V1 analyzed by fluorescent *in situ* hybridization (FISH) at P21 (see Figure S2 for other ages). E. Transcriptomic similarity is used to identify temporal associations among V1 glutamatergic neuron types across ages. Shown is a Sankey diagram, where nodes denote individual V1 glutamatergic neuron types at each age (as in Figure 1D), and edges are colored based on transcriptomic correspondence, determined using a supervised classification approach (see **Methods** for explanation). Darker colors indicate higher correspondence. F. Adjusted Rand Index (ARI) values quantifying temporal correspondence of glutamatergic types between each pair of consecutive ages based on transcriptomic similarity. Individual bars denote layers. ARI ranges from 0 (no correspondence) to 1 (perfect correspondence). Bar heights, mean ARI computed across pairs of consecutive ages; error bars, standard deviation; ***, *P*<0.0001 (one-way ANOVA) for layers 2/3 and 4 against layers 5 and 6. G. Types in L2/3 and L4, but not L5 and L6, are sensitive to changes in clustering resolution. Glutamatergic neurons at each age are re-clustered at different values of the resolution parameter (x-axis), and the results are compared with the base case corresponding to resolution = 1 (**Methods**). Line plots show mean ARI values for each layer (colors), while error bars denote standard deviation across different ages.

The relative proportions of most neuronal subclasses were stable over time (Figures 2C and S1L), although proportions of non-neuronal subclasses varied (Figure S1M). This suggests that the neuronal subclass composition of V1 is established before P8, our earliest time point. We also identified novel subclass-specific markers (Figures 2B and S2A-E). This included *Ccbe1* (collagen and calcium-binding EGF domain-containing protein 1), which is specific for L2/3 glutamatergic neurons throughout development (Figures 2D and S2A-C).

We performed dimensionality reduction and clustering for each class at each age separately. We henceforth refer to transcriptomically distinct clusters as types. The eight glutamatergic subclasses separated into 14-16 types, the six GABAergic subclasses separated into 14-15 types, and the six non-neuronal subclasses separated into 9-11 types depending upon age (Figure 1D) (**Methods**). Post-hoc differential expression analysis identified robust cell type-specific markers at each age (Figures S3A-C).

### Transcriptomic identities of L2/3 and 4 neuron types are established after eye opening

While the number of cell types within each class was similar at each age, it was not immediately clear how types identified at different ages were related to each other. Using transcriptomic similarity as a proxy for temporal relationships, we tracked the postnatal maturation of types within each class using a supervised classification framework (**Methods**). We observed striking subclass-specific differences in the maturation of glutamatergic neuron types (Figure 2E). L5, and to a slightly lesser extent L6, neuron types tightly corresponded throughout the time course, indicating that these types are established prior to eye-opening, and maintained. Conversely, upper-layer neuron types (L2/3 and L4) exhibited poor correspondences, suggesting gradual specification. Within L2/3, two neuron types at P8 and P14 matured into three types after eye-opening. By contrast, differences in the maturational patterns of GABAergic and non-neuronal subclasses were less pronounced (Figures S2F-I**, Methods**).

These subclass-specific differences in the timing of glutamatergic neuron type development are supported by five quantitative observations: (**1**) L5/6 types at different ages could be related in a 1:1 fashion with each other while L2/3/4 types could not be. These differences were based on the Adjusted Rand index (ARI), a measure of transcriptomic correspondence between two sets of clusters (Figure 2F). Furthermore, the clustering results for L2/3 and L4 were more sensitive (*P<0.0001,* one-way ANOVA) to changes in the resolution parameter than for L5 and L6 (Figure 2G); (**2**) The transcriptomic separation among L2/3 and L4 types was lower than that among L5 and L6 types, GABAergic types, and non-neuronal types at all ages (Figures S2J-L); (**3**) Differentially expressed genes that distinguished L2/3 and L4 neuron types varied with age, whereas those that defined L5 and L6 neuron types were stable (Figures S3D-G); (**4**) In a statistical test to identify temporally differentially expressed (tDE) genes in each layer (see **Methods**), L2/3 and 4 contained twice as many tDE genes as L5 and 6 (Figure S3H); and (**5**) The relative frequency of L2/3 and L4 types varied over time (see below). By contrast, the relative proportions of the ten L5 and L6 types, the smallest of which was present at an overall frequency of 1%, were stable throughout the time course. Together, these results suggest that within glutamatergic neurons of V1, transcriptomic specification of types within upper-layer subclasses (L2/3 and L4) occurs later than types in lower-layer subclasses (L5 and L6).

### L2/3 neuron types are spatially segregated

We classified L2/3 glutamatergic neurons into three types (A, B, and C) beginning at P17, the first time point assessed after the onset of vision at P14 (Figure 3A). These were visualized in tissue using *in situ* hybridization for marker genes *Cdh13*, *Trpc6*, and *Chrm2* for types L2/3_A, L2/3_B, and L2/3_C, respectively (Figures 3B-D). Cells expressing the three transcripts were organized into sublayers that became more pronounced with age: L2/3_A close to the pia, L2/3_C bordering L4, and L2/3_B in between (Figures 3D, E). At the boundaries of these sublayers, cells co-expressed more than one type-specific marker, indicating a lack of discrete, sharp boundaries, and mirroring the continuous transcriptomic arrangement observed *in silico* (see below).

**Figure 3.**
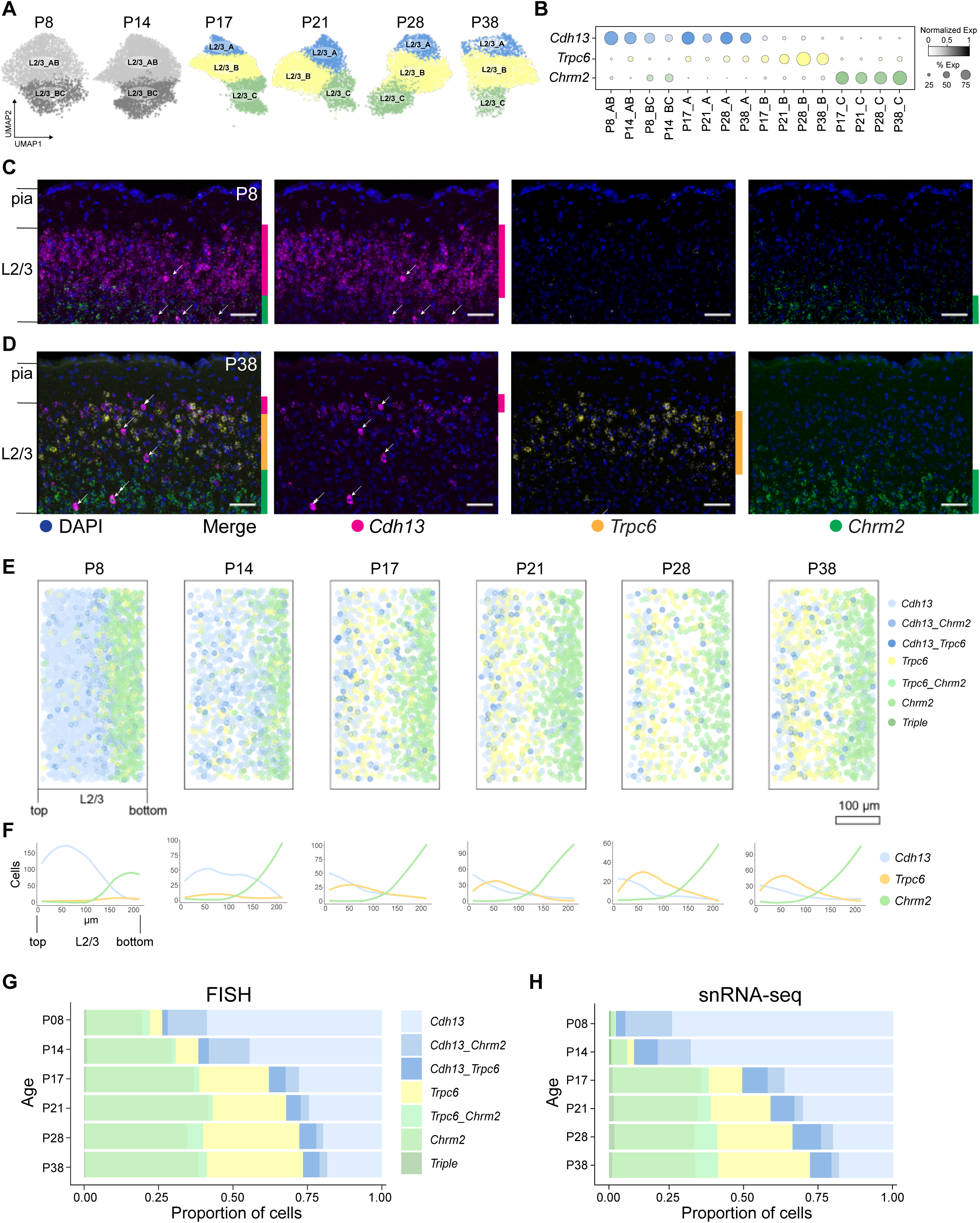
Anatomical and transcriptomic maturation of L2/3 glutamatergic neuron types. A. UMAP visualization of L2/3 glutamatergic neuron types across ages. B. Dot plot showing expression patterns of L2/3 type-specific genes (rows and colors) across L2/3 neuron types arranged by age (columns). Dot size represents the proportion of cells with non-zero transcript counts, and shading depicts log-transformed expression levels. C. FISH images showing type markers *Cdh13, Trpc6,* and *Chrm2* within L2/3 at P8. Vertical bars on the right of each image indicate sublayers expressing the indicated markers. Arrows, inhibitory neurons expressing *Cdh13*. Scale bar, 50 µm. D. Same as C, at P38. Arrows, inhibitory neurons expressing *Cdh13*. Scale bar, 50 µm. E. Pseudo-colored representation of *Cdh13, Trpc6,* and *Chrm2* expression in L2/3 cells across the six ages. Cells are grouped based on their expression levels of one or more of these markers (see legend on the right; **methods**). Each panel is an overlay of five or six images of V1 from three mice. Pial to ventricular axis is oriented horizontally from left to right within each panel. Total number of cells analyzed: P8, 2324; P14, 1142; P17, 1036; P21, 1038; P28, 653; and P38, 1034. Scale bar, 100 µm. F. Line tracings quantifying the number of cells per bin at each position along the pial to ventricular axis corresponding to panel E. 0 on the x-axis is the region of L2/3 closest to pia. 14 bins were used over the depth of L2/3. G. Relative proportions of cells within each expression group defined in panel E quantified using FISH data. H. Relative proportions of L2/3 cells within each expression group defined in panel E quantified using snRNA-seq data.

Prior to the onset of vision (P8 and P14), however, only two transcriptomic types were resolved. We denote these AB and BC. AB and BC were organized as two sublayers based on their differential expression of *Cdh13* and *Chrm2* (Figures 3C and S4A-E), with cells at the border co-expressing the two markers. In contrast, the B marker *Trpc6* was weakly expressed in cells scattered throughout L2/3 at these early stages (Figures 3B-C, E-H, and S4C, E). Multiple A-, B-, and C-specific markers were not expressed before P14 and only appeared at later stages (Figure S3D). Thus, we infer that the L2/3 glutamatergic types A, B, and C arise from AB and BC types following the onset of vision (Figure 2E and **Methods**).

### Vision is necessary for establishing and maintaining L2/3 neuron type identity

The emergence of three L2/3 neuron types following eye-opening prompted us to explore the role of vision in defining cell types. It is well established that vision is required for the development of cortical circuitry during the critical period for ocular dominance plasticity (P21-P38) ^14,17^. We used snRNA-seq to profile V1 in animals that were dark-reared from P21 to P28 and P21 to P38. For brevity, these experiments are referred to as P28DR and P38DR, respectively (DR = dark rearing). We also profiled animals that were exposed to 8 hours of ambient light after dark-rearing from P21-P28 to assess the impact of visual stimulation following prolonged deprivation (Figure 4A). We refer to this experiment as P28DL (DL = dark-light). In total, we recovered 77,150 high-quality nuclei across these three experiments and identified classes, subclasses, and types using the same computational pipeline applied to the normally reared (NR) samples (Figure 4B and **Methods**).

**Figure 4.**
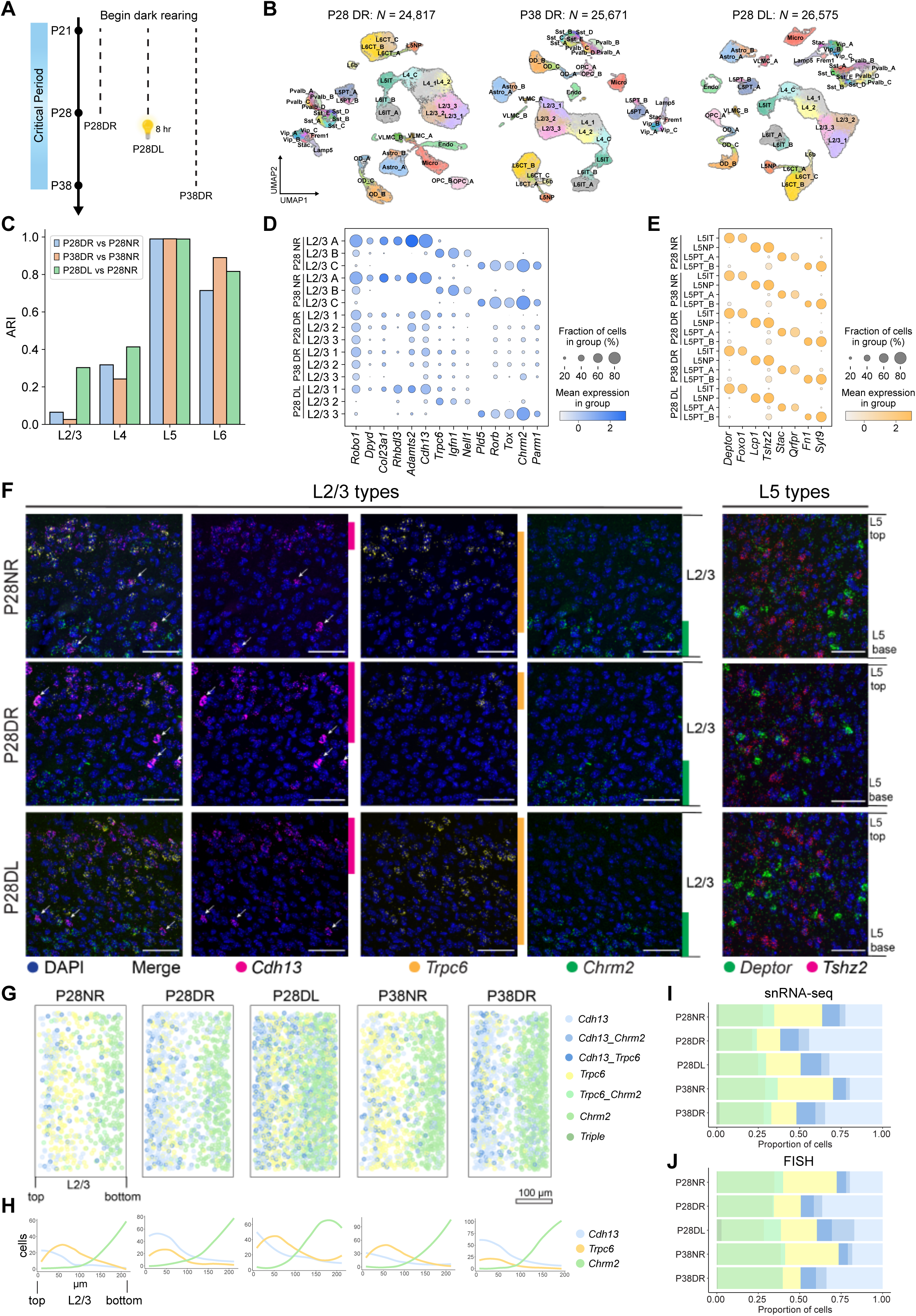
Visual experience is required to maintain L2/3 glutamatergic neuron types. A. Schematic of experiments to probe the influence of visual input on V1 transcriptomic identities. Data were collected from three rearing conditions: Dark-reared between P21-P28 and snRNA-seq profiling (P28DR), dark-reared between P21-P38 and snRNA-seq profiling (P38DR), and dark-reared between P21-P28 followed by 8h of ambient light stimulation and snRNA-seq profiling (P28DL). B. UMAP visualization of transcriptomic diversity in P28DR (*left*), P38DR (*middle*), and P28DL (*right*). Clusters that match 1:1 to normally-reared (NR) types in Figure 1D are labeled accordingly. This was not possible for all L2/3 and two L4 clusters, which correspond poorly to NR types. We therefore provisionally labeled these clusters L2/3_1, L2/3_2, L2/3_3, L4_1, and L4_2. C. Bar plot showing the Adjusted Rand Index (ARI) quantifying transcriptomic similarity within each layer (x-axis) between glutamatergic clusters observed in dark-reared mice and types observed in normally-reared (NR) mice. Colors correspond to comparisons: P28DR vs. P28NR, P38DR vs. P38NR, and P28DL vs. P28NR. D. Expression of L2/3 type markers (columns) in NR, DR, and DL types and clusters (rows) at P28 and P38. E. Same as panel D for L5. DR and DL clusters are labeled based on their tight transcriptomic correspondence with NR types (Figure S5F, G). F. FISH images showing expression of L2/3 and L5 type markers in NR (*top row*), DR (*middle row*), and DL (*bottom row*) at P28. *Deptor* is a marker for L5IT and *Tshz2* is a marker for L5NP. Arrows, inhibitory neurons expressing *Cdh13*. Scale bar, 50 µm. G. Pseudo-colored representation of *Cdh13, Trpc6,* and *Chrm2* expression in L2/3 cells in NR, DR, and DL mice at P28 and P38 (see legend on the right). Each plot is an overlay of 5-6 images of V1 from three mice. Pial to ventricular axis is oriented horizontally from left to right within each panel. Total number of cells analyzed: P28NR, 653; P28DR, 989; P28DL, 1732; P38NR, 1034; and P38DR, 1177). Scale bar, 100 µm. H. Line tracings quantifying the number of cells per bin at each position along the pial to ventricular axis corresponding to panel G. 0 on the x-axis is the region of L2/3 closest to pia. 14 bins were used over the depth of L2/3. I. Relative proportions of L2/3 cells within each expression group defined in panel G quantified using snRNA-seq data J. Relative proportions of cells within each expression group defined in panel G quantified using FISH data.

We performed three computational analyses to probe the effect of visual deprivation (DR and DL) on the transcriptomic patterns observed in normally reared (NR) mice. First, we compared the overall transcriptional profiles of cell types across the three conditions. We found that dark rearing disrupted the type identities of L2/3 and L4, but not L5 and L6, glutamatergic neurons (Figures 4C and S5F). Furthermore, dark rearing neither altered the gene expression patterns that defined subclasses nor those defining GABAergic and non-neuronal cell types (Figures S5A-C). Second, the cell type markers identified in NR mice were disrupted by dark rearing in L2/3, and slightly in L4, but not L5 and 6 (Figures 4D-E and S5D-E). Third, we probed the effect of visual deprivation on type-specific genes within each layer. While the signatures within all four layers were different when comparing DR to NR, the effect was most dramatic for L2/3 (Figure S6H). Thus, vision selectively influences transcriptomic profiles of upper-layer glutamatergic cell types.

The effect of dark rearing was particularly striking in L2/3. The L2/3 clusters observed in dark-reared mice poorly resembled the three types in normally reared animals, and the expression patterns of cell type-specific marker genes were disrupted (Figure 4C, D). By contrast to the three sublayers highlighted by *Cdh13, Trpc6,* and *Chrm2* expression in L2/3 in normally reared mice, only two sublayers were observed in dark-reared mice. Notably, there was a sharp decrease in *Trpc6*-expressing cells (Figure 4F-J), consistent with snRNA-seq data (Figure 4D). This was not simply a loss of one cell type, however, but a global disruption of gene expression patterns throughout L2/3 (see below, Figure 6). The two-layered pattern was more prominent in dark-reared animals at P38 compared to P28 (Figures 4G-I and S5H-I). Thus, in the absence of vision, the expression patterns of these markers were similar to those prior to the onset of vision (see panels P8 and P14 in Figure 3E).

The loss of cell type identity in animals deprived of light during the first half of the critical period was partially reversible. L2/3 transcriptomic clusters in mice exposed to 8 hours of ambient light after dark rearing between P21-P28 showed a marked recovery of gene expression patterns observed in normally reared animals (Figures 4C-D and S5G, S6H). In addition, the layered arrangement of *Cdh13-*, *Trpc6-* and *Chrm2*-expressing cells in these animals was also shifted towards that observed in normally reared animals (Figures 4F-J and S5H-I). These results demonstrate that vision is needed to maintain the transcriptomic and spatial identities of L2/3 cell types.

As the spatial expression of cell type markers in the absence of vision and at eye opening were similar, we set out to assess whether vision was not only necessary to maintain cell types, but also required for their establishment. To test this, we dark-reared mice from P8 to P17 (Figure 5A) and assessed the expression patterns of *Cdh13*, *Trpc6* and *Chrm2* in tissue sections. These mice had two, instead of three, sublayers within L2/3, similar to P8 and P14 normally reared animals (Figures 5B-D). These changes included a dramatic reduction in *Trpc6-*expressing cells and an increase in *Cdh13* expression, which was accompanied by an expansion in its expression domain towards the middle sublayer in mice with no visual experience (Figure 5E). This was not a general effect on glutamatergic cell types, as the relative proportions of L5 neuron types were insensitive to changes in visual experience (Figure 5F). In summary, these results show that vision acts selectively in L2/3 to establish and maintain cell types.

**Figure 5.**
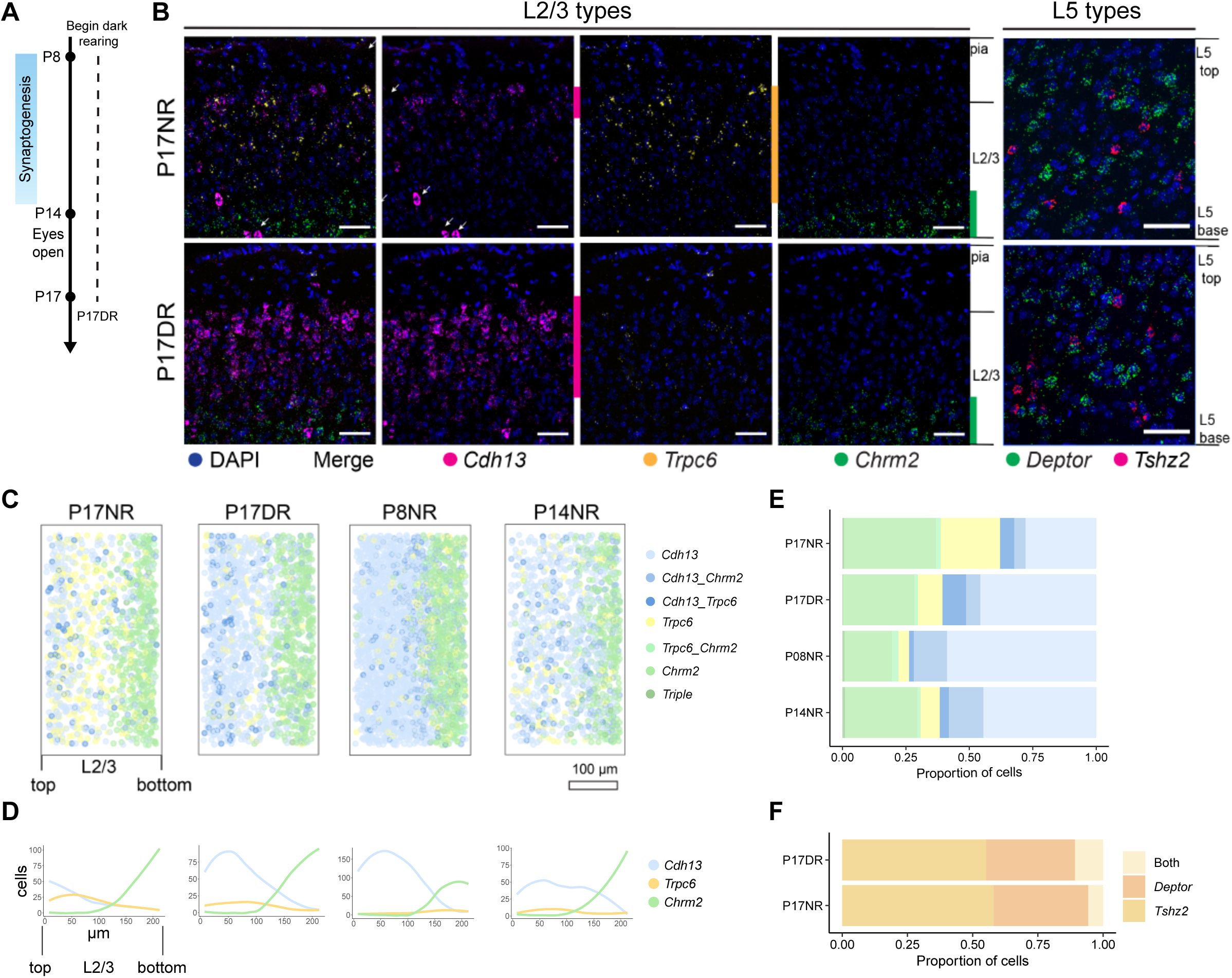
Vision is required to establish L2/3 glutamatergic neuron types. A. Schematic of experiments to probe the role of vision in establishing three L2/3 glutamatergic types. B. FISH images showing expression of L2/3 and L5 type markers in normally-reared (NR) and dark reared (DR) mice at P17. Arrows, inhibitory neurons expressing *Cdh13*. Scale bar, 50 µm. *Deptor* is a marker for L5IT and *Tshz2* is a marker for L5NP. C. Pseudo-colored representation of *Cdh13, Trpc6,* and *Chrm2* expression in L2/3 cells (see legend on the right). Each plot is an overlay of 6 images of V1 from three mice. Total number of cells analyzed: P17NR, 1036; P17DR, 1411. D. Line tracings quantifying number of cells per bin at each position along the pial to ventricular axis corresponding to panel C. 0 on the x axis is the region of L2/3 closest to pia. 14 bins were used over the depth of L2/3. E. Relative proportions of cells within each expression group defined in panel C quantified using FISH data. F. Relative proportions of two L5 cell types, L5IT (*Deptor*^+^) and L5NP (*Tshz2*^+^) in normal and dark reared mice at P17. Total number of cells analyzed: P17NR, 1676; P17DR, 1544.

### Continuous variation of L2/3 neuron types and gene expression gradients are shaped by vision

The sublayers corresponding to types A, B, and C in L2/3 were partially overlapping, mirroring the continuous arrangement of their transcriptomes (see Figures 3A, E). Consistent with this continuous arrangement, more than 70% of the 285 differentially expressed genes among the L2/3 types in normally reared mice exhibited graded, rather than digital, differences (Figures 6A and S6A-H). In dark-reared mice, these genes were no longer expressed in a graded fashion between the L2/3 clusters, although their overall (i.e., bulk) expression levels, for all but a few genes, were unaltered (Figures S6C). These gradients were partially recovered by brief restoration of normal visual experience to dark reared animals during the critical period (Figure 6A). Thus, vision selectively regulates gene expression in a sublayer-specific fashion, contributing to the continuous variation of L2/3 cell types.

**Figure 6.**
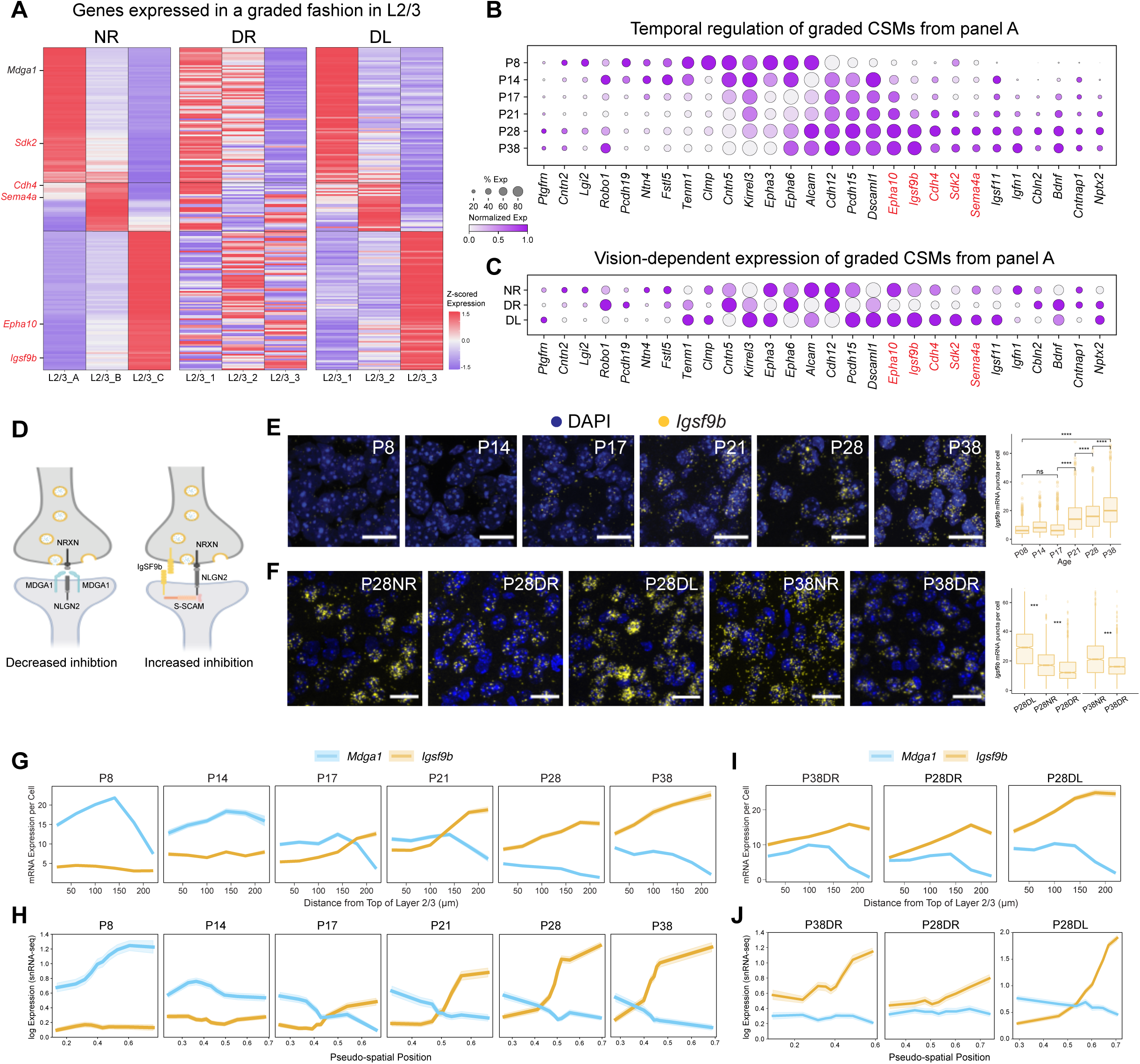
Continuous variation of L2/3 neuron types and vision-dependent gene gradients implicated in wiring. A. Heatmap of L2/3 type-specific genes showing graded expression in normally-reared mice (NR, *left panel*). This is disrupted in dark-reared mice (DR, *middle panel*) and partially recovered by exposing these DR mice to ambient light for eight hrs (DL, *right panel*). For the full set of L2/3 type-specific genes grouped by expression pattern, see Figure S6A. Genes that satisfy the criteria outlined in panels B and C (see text) are indicated in red lettering on the left. B. Dot plot showing the temporal regulation of cell surface molecules (CSMs) found in panel A. Highlighted in red are genes that are upregulated during the classical critical period (P21-P38), and as shown in panel C, downregulated in response to dark-rearing (DR) and upregulated in response to DL rearing. Expression levels shown for L2/3 neurons. C. Same illustration as panel B across the conditions P28NR, P28DR, and P28DL. D. Schematic of MDGA1 and IGSF9B interactions with NLGN2 at synapses. MDGA1 prevents NLGN2 interaction with NRXN presynaptically. IGSF9B binds homophilically across the synapse and interacts with S-SCAM postsynaptically to stabilize NLGN2 interaction with NRXN. E. (*Left*) FISH images showing the expression of *Igsf9b* mRNA over time in V1. Three animals per time point, six images per animal. Scale bar, 20 μm. (*Right*) Box plot quantifying expression. Wilcoxon Rank Sum Test, **** p <0.0001. Number of cells quantified: P8,1191; P14,1011; P17, 1389; P21, 1729; P28, 1277; and P38, 1588. F. (*Left*) FISH images showing that dark rearing decreases *Igsf9b* expression in L2/3, and exposure to ambient for eight hrs of light restores expression. Scale bar, 50 µm. (*Right*) Box plot quantifying expression. Three animals imaged per age and condition combination. Number of cells quantified: P28NR, 1290 cells; P28DL, 1506 cells; P28DR, 1521 cells; P38NR, 1629 cells; and P38DR, 1885 cells. Quantified at 40X. Wilcoxon Rank Sum Test, *** p <0.001. G. FISH quantification of average *Mdga1* and *Igsf9b* expression (y-axis) in glutamatergic cells as a function of distance from the top of L2/3 (x-axis). Shaded ribbons represent standard error of the mean. Panels represent different ages. Cell numbers at each age: P8, 2204; P14, 928; P17, 1037; P21: 1183; P28, 719; and P38: 942 cells. Data collected from three to four animals at each age. H. Reconstruction of *Mdga1* and *Igsf9b* expression levels averaged across cells based on their inferred L2/3 pseudo-spatial locations in gene expression space (see **Methods** for details). Shaded ribbons represent standard deviation. I. Same as panel G for P38DR, P28DR, and P28DL. Cell numbers: P38DR, 719; P28DR, 1061; and P28DL, 1053 cells. Data collected from three animals at each time point J. Same as panel H for P38DR, P28DR, and P28DL. Note difference in scale for P28DL to capture the extent of increase in *Igfs9b* expression.

We hypothesized that graded genes which are temporally regulated and expressed in a vision dependent manner could be associated with functional changes in L2/3 during the critical period. Several genes fit this description, including cell surface molecules (CSMs) and transcription factors (TFs) (Figures 6B, C and S6I, J). Among these were cell surface and secreted proteins previously shown to be involved in the development of neural circuits, including proteins regulating cell recognition (e.g., *Kirrel3*, *Sdk2*) and synaptic adhesion (e.g., *Tenm1* and *Cbln2*) (Figures 6B, C). To identify candidate cell surface proteins from this set that may contribute to vision-dependent changes in circuitry during the critical period, we selected genes that satisfied three criteria across all L2/3 glutamatergic types: 1. Selective upregulation during the critical period; 2. Downregulation in DR animals; and 3. Upregulation in DR animals in response to eight hours of ambient light at P28. Five genes (*Epha10, Igsf9b*, *Cdh4*, *Sdk2*, and *Sema4a)* satisfied all three criteria, with the expression levels and dynamics of *Igsf9b* being the most robust (Figures 6E, F).

### *Igsf9b* knock-out alters inhibitory synapses in L2/3

IGSF9B is a homophilic cell adhesion molecule of the immunoglobulin superfamily that promotes Neuroligin2 (NLGN2)-dependent inhibitory synapse formation ^33,34^ (Figure 6D). This protein is of particular interest as inhibition plays an important role in regulating V1 circuitry during the critical period ^13^. We assessed the spatial distribution of *Igsf9b* transcripts in L2/3 at different times in development using FISH (Figure 6G) and *in silico* by regarding the transcriptomic positions of L2/3 neurons in gene expression space as “pseudo” spatial coordinates (Figure 6H**; Methods**). *Igsf9b* levels were low prior to eye opening and increased during the critical period in a graded manner favoring increased expression deeper into L2/3. Sensory activity further modulated the expression level and lamination of *Igsf9b*. Dark-rearing during the critical period decreased *Igsf9b* expression and slightly disrupted its graded expression in L2/3 neurons (Figures 6F, I, J). These effects were reversed and *Igsf9b* expression levels were upregulated when dark-reared animals were exposed to ambient light for 8 hrs. Given the graded expression of *Igsf9b* across sublaminae increasing from upper to lower layers, it was particularly intriguing that a second gene encoding another Ig superfamily protein*, Mdga1*, a negative regulator of NLGN2 (Figure 6D), was expressed in a graded and opposite spatial pattern to *Igsf9b* (Figure 6G, H; Figure S6F). Together, the spatiotemporal dynamics of *Igsf9b* and *Mdga1* expression after eye opening form a gradient of inhibitory synapse potential along the pial-ventricular axis of L2/3, with lower sublayers exhibiting increased inhibition.

To explore the role of *Igsf9b* in the development of inhibitory synapses in L2/3, we examined the expression of five markers of inhibitory synapses in wild-type (WT) and *Igsf9b* knock-out (KO) mice. These markers included three postsynaptic proteins (Gamma-aminobutyric acid Type A receptor subunit alpha 1 (GABRA1), Neurolign2 (NLGN2), and Gephryin (GPHN)) and two presynaptic proteins (the presynaptic vesicular GABA transporter (VGAT) and the enzyme glutamic acid decarboxylase (GAD65)) (Figure S7A). Expression levels of the postsynaptic markers GABRA1 and NGLN2 were significantly decreased in P37 KO mice relative to WT littermates (Figure S7B), although GPHN remained unchanged (not shown). By contrast, there was an increase in the levels of presynaptic markers GAD65 and VGAT (Figure S7B); this increase may reflect a homeostatic response to the changes reflected in the decrease in postsynaptic markers. Consistent with our finding that expression of *Igsf9b* in L2/3 increases with depth, these phenotypes were more pronounced toward the bottom of L2/3 (Figure S7D-G; see Figure 6G, H). By contrast, excitatory synapse markers were unaffected in KO mice (Figure S7C). Thus, loss of *Igsf9b* specifically affects inhibitory synapses in a graded fashion along the L2/3 pial-ventricular axis.

### *Igsf9b* regulates vision-dependent maturation of binocular circuitry

A defining feature of the critical period in V1 is the vision-dependent maturation of binocular neurons, which are required for depth perception, also known as stereopsis ^35^. To mediate stereopsis, these neurons must selectively respond to the same kind of visual information from both eyes (binocular matching). Although binocular neurons can be detected shortly after eye opening, they exhibit poor matching at early stages. Visual experience throughout the critical period (P21-P36) is required to replace these early binocular neurons with new binocular neurons that are properly matched ^15,16^.

We examined whether *Igsf9b* is required for the normal development of binocular responses in L2/3 using *in vivo* 2-photon calcium imaging in binocular V1 (Figure S7H, I). This region of V1 comprises not only neurons responsive to both eyes (i.e., binocular cells), but also monocular neurons responsive to stimuli presented to the ipsilateral or contralateral eye only. We measured responses of thousands of excitatory neurons to stimulation of each eye at P21 and P36 in normally reared WT and *Igsf9b* KO mice (Figure 7A; see **Methods**). These results were also compared to those from GCaMP6s transgenic mice that were normally reared (NR) or dark-reared (DR) during the critical period ^15^(Figure 7A). Neuronal responses were measured at three depths, spanning the top (type A), middle (type B), and bottom (type C) sublayers of L2/3, corresponding to the regions of low, intermediate, and high *Igsf9b* expression, respectively (Figure 7B-D, Figure S7J-L; see **Methods**). At P21, the proportion of binocular neurons and their orientation matching in KO mice were indistinguishable from those in WT (Figure 7E). This is consistent with the low expression of *Igsf9b* in L2/3 before P21 (Figure 7A, see Figure 6). At P36, after closure of the critical period—when *Igsf9b* expression would normally have increased—only about half the normal number of binocular neurons were observed in KO mice, and these few neurons displayed poor binocular matching (Figure 7F). This phenotype resembles that observed in DR mice (Figure 7G).

**Figure 7.**
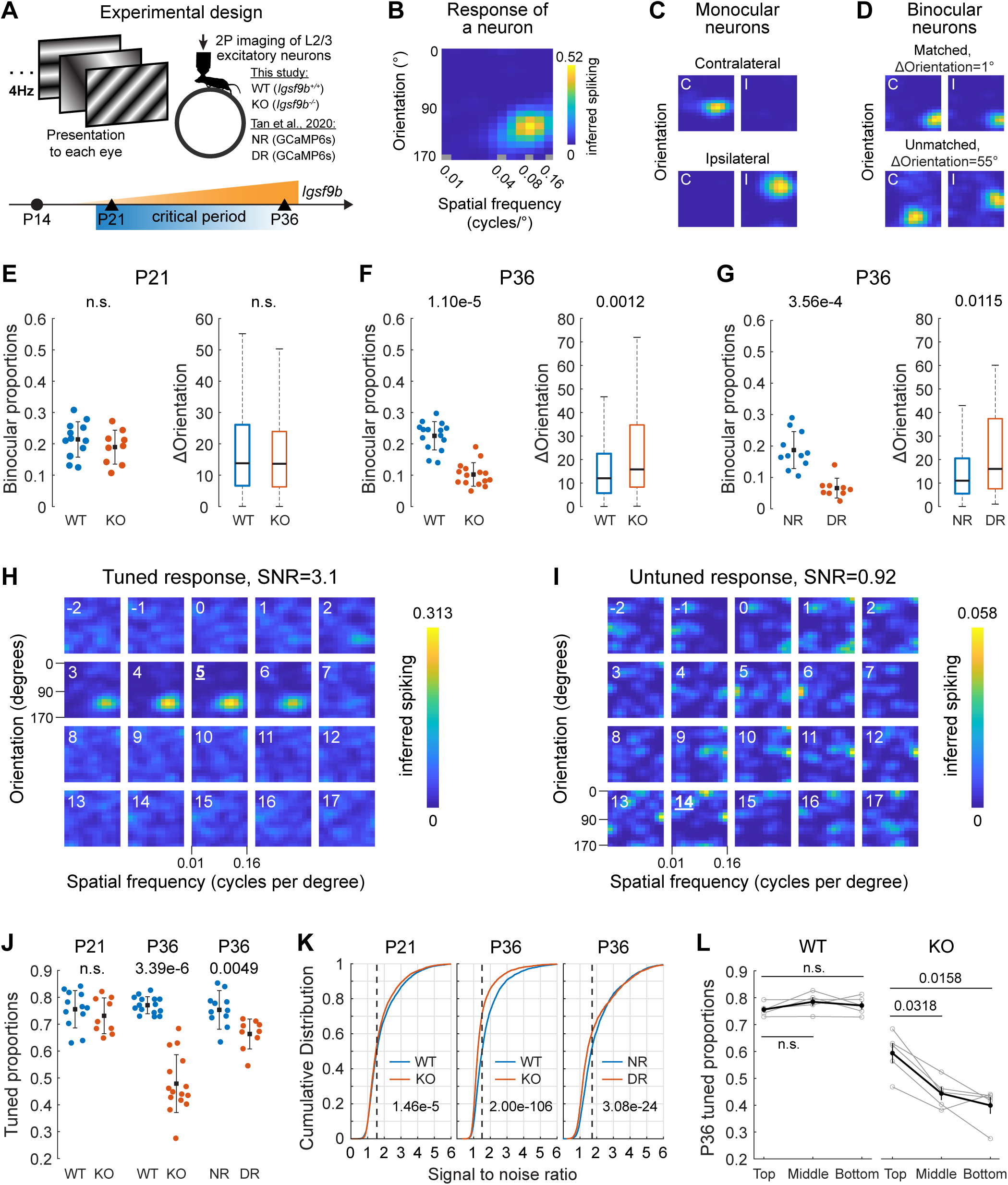
Igsf9B is required for vision-dependent maturation of binocular neurons in V1B L2/3. A. Experimental set up for functional analysis on *Igsf9b* by imaging visual responses of layer 2/3 excitatory neurons in the binocular region of V1 (V1b). *Top panel*, schematic of 2-photon (2P) calcium imaging using a battery of sinusoidal gratings sequentially presented to mouse at 4 Hz. Visual stimuli were presented to each eye separately. The head-fixed mouse was awake on a running wheel. Mice used in this study are WT (*Igsf9B^+/+^*) and KO (*Igsf9B^-/-^*) expressing AAV encoded jGCaMP7f. Panels G, J, K include our unpublished results from normally reared (NR) and dark reared (DR) (dark reared from P22-P36) transgenic mice carrying GCaMP6s expressed in excitatory neurons (from Tan et al., 2020). *Bottom panel*, schematic of the experiment design in this study. WT and KO mice were imaged using 2P microscope at P21 and P36, the onset and closure of the classical critical period. Orange is a representation of changes *Igsf9B* expression in layer 2/3 as a function of age. B. Tuning kernel showing response of a single neuron (see Figure S7L) to the contralateral eye. The colors represent response strength (color bar, right) as a function of stimulus orientations (Y-axis) and spatial frequency (log scale; X-axis). C. Tuning kernels to the contralateral (C) or ipsilateral eye (I) from a monocular contralateral neuron (top) and a monocular ipsilateral neuron (bottom). Kernels for each neuron were normalized to the peak inferred spiking of the neuron. D. As in C, but for a matched binocular neuron (top) and an unmatched binocular neuron (bottom). The difference in orientation preference to one or the other eye, or ΔOrientation, is shown above the kernels for each neuron. E. Left panel, proportions of binocular neurons in WT and KO mice at P21. Each point is from a single imaging plane. Mean and standard deviation, black dots and lines. Mann-Whitney U test. Right panel, boxplots of ΔOrientation of binocular neurons in WT (4 mice, 761 cells) and KO (3 mice, 619 cells) mice at P21. Black horizontal line, median; box, quartiles with whiskers extending to 2.698σ. Mann-Whitney U test. Note the absence of phenotype in binocular neurons at P21. F. As in E but for binocular neurons at P36. WT, 5 mice, 602 cells; KO, 5 mice, 269 cells. G. As in F but for binocular neurons in NR (4 mice, 339 cells) and DR (3 mice, 78 cells) mice. Note the P36 phenotypes in KO and DR mice were similar. The difference in proportion between the WT (panel F) and NR (panel G) likely reflects differences in genetic background or experimental design (i.e., viral versus transgenic expression of GCaMP or differences between GCaMP6s and jGCaMP7f). Analysis of tuned neurons, including monocular and binocular neurons. H. Example of a tuned cell from a WT mouse at P21. Inferred spiking as a function of imaging frames for a neuron with a tuned response. Numbers at the top left indicate imaging frames relative to stimulus onset. For this neuron, the SNR is 3.1, and peak response occurred 5 imaging frames or 323 ms after onset of its optimal stimuli, consistent with the kinetics of jGCaMP7f. I. As in H but for an untuned cell in the same mouse at P21. For this neuron, the SNR is 0.92, and peak response occurred 14 imaging frames or 903 ms after stimulus onset, lagging far behind the normal temporal dynamics of jGCaMP7f. J. Proportions of tuned neurons in WT and KO mice at P21 (left two) and P36 (middle two), and in NR and DR mice at P36 (right two). Each point is from a single imaging plane. Mean and standard deviation, black dots and lines. Mann-Whitney U test. K. Left panel, cumulative distribution of signal to noise ratio (SNR) to either eye of all imaged neurons at P21 in WT (4 mice, 3436 neurons ) or KO (3 mice, 3457 neurons) mice. Dashed vertical line marks the SNR threshold for visually evoked responses (see **Methods** for details). P-value from two-sample Kolmogorov-Smirnov test was shown in the plot. Middle panel, as in the left, but for mice at P36 in WT (5 mice, 2698 neurons) and KO (5 mice, 2699 neurons). Right panel, as in the middle, but for neurons in NR (4 mice, 1905 neurons) and DR (3 mice, 1188 neurons) mice. In all cases, the response of a neuron to each eye was measured separately. L. Proportions of tuned neurons as a function of depth in V1B layer 2/3 in WT and KO mice at P36. Top, middle, and bottom indicate the three imaging planes covering the corresponding sub-laminae within layer 2/3 in each mouse. Each gray line represents a mouse. Mean and standard error of the mean were shown as black dots and vertical lines. Mann-Whitney U test with Bonferroni correction.

We also noted a marked decrease in the proportion of tuned cells in KO mice. Untuned cells were active, but by contrast to tuned cells, they were not responsive to specific visual stimuli in a time-locked fashion (Figure 7H, I). While visual deprivation decreased the proportion of visually responsive neurons from 75% (NR) to 66% (DR), *Igsf9b* KO decreased the proportion of responsive neurons to 47% by P36 (Figure 7J). This reduction in proportion of tuned neurons in KO mice from P21 to P36 correlated with a marked reduction in the signal to noise ratios (SNRs) of neuronal responses at critical period closure (Figure 7K, left and middle panels). The extent of SNR impairment in KO was more severe than in DR mice (Figure 7K, compare middle and right panels). Notably, the severity of reduction of tuned neurons in KO increased with depth (Figure 7L), mirroring the graded expression of *Igsf9b* in L2/3 types A, B, and C along the pial-ventricular axis in normally reared WT mice (see Figure 6). Taken together, these findings establish that *Igsf9b* regulates the vision-dependent maturation of L2/3 excitatory neurons in a graded fashion along L2/3 pial-ventricular axis.

## DISCUSSION

Critical periods define windows of postnatal development where neural circuitry is particularly sensitive to experience. Here we sought to gain insight into how experience influences circuitry during this period at the level of cell types using single-nucleus sequencing of primary visual cortex. We showed that, unexpectedly, vision regulates cell type specification to establish a continuum of L2/3 glutamatergic cell types arranged in sublayers along the pial-ventricular axis. We show that genes encoding cell surface proteins that regulate wiring are also expressed in this graded pattern. This raises the prospect that these proteins directly regulate changes in circuitry during this period. In support of this idea, we demonstrate that one of these, IGSF9b, an adhesion molecule at inhibitory synapses, regulates the development of binocular responses in L2/3 glutamatergic neurons in a graded fashion during the critical period.

### A postnatal developmental atlas of mouse V1

To study vision-dependent cortical development at the level of cell types, we generated a developmental atlas of mouse V1 comprising over 220,000 nuclear transcriptomes spanning six postnatal ages and three light-rearing conditions. Several features of this dataset enabled us to identify robust and reproducible biological signals. First, we identified a similar number of transcriptomic clusters at all six ages, which were collected and processed separately. For all clusters, transcriptomic identities and relative proportions were comparable between independent samples, consistent with these being bona fide cell types. Second, computational inference of transcriptomic maturation showed that the GABAergic, deep-layer glutamatergic, and non-neuronal cell types were present prior to eye opening and remained largely unchanged through the critical period, whether animals were reared in a normal dark/light cycle or in the dark. Third, these stable cell types served as important “negative controls” that enabled us to identify the minority of cell types among the upper layer glutamatergic neurons that were specified following eye opening, and whose transcriptomic identities were profoundly influenced by vision. Fourth, we identified novel cell type markers that enabled us to uncover the arrangement of L2/3 cell types in sublayers (see Figures 3B-D). And finally, the developmental atlas served as a foundation to investigate vision-dependent functional maturation of V1 at the resolution of cell types and molecules.

### The establishment and maintenance of L2/3 neuron types require vision

Both transcriptomic cell type identity and cell function emerge gradually in L2/3 glutamatergic neurons after eye opening. In previous studies, we showed that there are few binocular neurons in L2/3 at eye opening. Their numbers increase over the next several days in a vision-dependent process ^16^. During the critical period, these binocular neurons, most of which are poorly tuned, are rendered monocular. In parallel, new binocular neurons are formed from the conversion of other well-tuned monocular neurons which gained matched responses to the other eye ^15^. It is through this exchange of neurons that well-tuned and matched binocular neurons emerge to give rise to mature binocular circuitry. These changes rely on vision during the critical period.

It is striking that the acquisition of transcriptomic cell type identity in L2/3 follows a similar time course. Although it is well known that neural activity regulates gene expression ^21,36,37^, our finding that vision establishes and maintains cell type identity broadens the role of neural activity in regulating cellular development. Transcriptomic changes could arise through changes in circuitry driven by activity. Alternatively, activity may drive cell type changes that, in turn, instruct changes in circuit organization. Further experiments will be necessary to distinguish between these and other mechanisms. Experience-dependent regulation of upper-layer cortical cell types may be a general principle underlying cortical development during critical periods.

### Continuous variation in L2/3 identity and sublayer arrangement

Although unsupervised clustering defined three predominant glutamatergic neuronal types in L2/3, the gene expression differences between them were graded, giving rise to continuous variation in transcriptomic identity (Figures 6A and S6A-B). This continuous variation *in silico* was seen as a spatially graded, sublayered arrangement in L2/3 via FISH. This is not a general feature of glutamatergic cell type specification, as glutamatergic cell type identity in L5, for example, is neither graded nor dependent upon vision. Continuous variation of cell type identity has been reported in other regions of the mammalian brain^38–41^. This contrasts the transcriptomic features of neurons in the fly brain and mammalian retina where cell types exhibit molecularly discrete identities, even for highly related neurons^42–44^.

That the molecular heterogeneity in L2/3 reflects functional differences is supported by a recent retrograde labeling analysis of adult V1, which identified transcriptional signatures of L2/3 glutamatergic neurons that project to the anterolateral (AL) and posteromedial (PM) higher visual areas (HVAs) ^45,46^. PM- and AL-projecting neurons localize in the upper and lower regions of L2/3 and express the markers *Cntn5/Grm1* and *Astn2/Kcnh5*, respectively (Figure S6D). In our data, these markers are expressed in a graded and opposite fashion along the pial-ventricular axis, suggesting that PM- and AL-projecting L2/3 neurons localize to the upper (type A) and lower (type C) sublayers, respectively (Figure S6E). Interestingly, in this same study, other L2/3 neurons found in a broad domain between the upper and lower layers targeted their axons preferentially to the lateromedial (LM) HVA. These cells occupy a region similar to cell type B in our studies.

L2/3 neurons form numerous “local” circuits that process diverse visual information, but these are yet to be defined at the cell type level ^47^. While a given excitatory neuron may participate in multiple circuits, there is evidence for synaptic specificity. For instance, neurons projecting to PM show a strong preference for connectivity with PM-projecting neurons over AL-projecting neurons ^45^. It is tempting to speculate that this functional segregation may, in part, be due to graded molecular differences between neurons uncovered in our transcriptomic analysis. These graded differences may also exist along other spatial axes to further subdivide sublayers into functional circuits.

### Experience-dependent cell type specification in L2/3

One of our main findings is that vision specifies L2/3 cell types in V1 and that these cell types are arranged in a sublayered fashion. Recent studies have reported the sublayered organization of cell types in L2/3 of the mouse motor cortex, in addition to the visual cortex ^48–53^, suggesting an analogous experience-dependent mechanism may be involved in patterning this region. Emerging transcriptomic, morphological, and physiological evidence of similar cell type continuums arranged in sublayers in L2/3 of the human cortex (Berg et al., 2021) raise the exciting possibility that experience-dependent cell-type specification may be a general principle of mammalian cortical development. A sublaminar organization of cell types may facilitate preferential local connectivity between glutamatergic neurons of the same type and would provide an efficient strategy to organize their axonal projections to the same higher visual areas and facilitate recurrent connections from these back to their appropriate sublayers ^5^.

### Igsf9b is a vision-dependent regulator of cortical circuitry

Patterns of experience-dependent activity may promote the expression of recognition molecules that regulate wiring. Indeed, different patterns of experience-independent activity have been shown to regulate the expression of cell-type specific wiring genes in the mouse olfactory system ^54^. Our identification of vision-regulated recognition molecules expressed in L2/3 neurons, and detailed genetic studies on Igs9b provide support for the view that experience-dependent regulation may also play a role in establishing cell-type specific wiring (Figures 7 and S7).

Analyses of V1 in mice that lack *Igsf9b* revealed changes in inhibitory, but not excitatory, synaptic markers. More significantly, *Igsf9b^-/-^* mice showed a significant decrease in the proportion of binocular neurons and the proportion of tuned neurons. In the case of the latter phenotype, the severity of the defect increased in deeper sublayers, where in wild type animals, *Igsf9b* expression is higher. A similar impact on the tuning of glutamatergic neurons, more broadly within L2/3, was observed in optogenetic experiments in which perisomatic PV inhibitory neuron activity was suppressed ^55^. Thus, IGSF9b may modulate PV inhibitory input onto L2/3 excitatory neurons during the critical period. These findings are consistent with previous studies demonstrating a role for IGSF9b in regulating inhibitory synapses.

In summary, our results raise the exciting possibility that experience-dependent cell type specification is a general phenomenon in mammalian brain development. Specific patterns of neural activity arising as a consequence of experience may lead to graded changes in the expression of genes which, in turn, alter neural circuits. Understanding how the interplay between circuit function, cell type specification, and experience sculpts circuitry will rely on integrating multiple levels of analysis from molecules to behavior.

## Supporting information

Supplemental Table 1, and will be used for the link to the file on the preprint site.

Supplemental Table 2, and will be used for the link to the file on the preprint site.

## ACKNOWLEDGEMENTS

The authors would like to thank Maria del Carmen Diaz de la Loza for illustrations. Select images were made using BioRender. We thank Xiang Li, Michael Mashock, and Xinmin Li from UCLA Technology Center for Genomics and Bioinformatics for assistance with scRNA-seq. We also thank Andrew Elkins and Drs. Damon Polioudakis and Dan Geschwind for advice and help at early stages in this project. We are grateful to Drs. Weize Hong, Jonathan Flint, Emile Marcus, Dario Ringach, and Joshua Sanes, and members of the Shekhar and Zipursky labs for critical feedback. We thank Drs. Nathan Gouwens, Clay Reid, Staci Sorensen, Bosiljka Tasic, Zizhen Yao, and Hongkui Zeng for helpful discussions and sharing unpublished data. This work was supported by the Stein Eye Institute EyeSTAR program (S.C.), NSF Graduate Research Fellowship (grant DGE1752814 to S.B.), an NIH grant (R00EY028625 to K.S.), a grant from the W.M. Keck Foundation to S.L.Z and J.T.T., and startup funds from the University of California, Berkeley (K.S). S.L.Z. is an investigator of the Howard Hughes Medical Institute.

## AUTHOR CONTRIBUTIONS

S.C., L.T., J.T.T., and S.L.Z conceived the project. S.C., L.T. and S.L.Z. designed snRNA-seq experiments. S.B. and K.S. conceived and designed the computational methods for transcriptomic analysis. S.C. carried out snRNA-seq and initial bioinformatic analysis. S.B. and S.S. performed the transcriptomic analysis. S.C. and V.X designed and carried out FISH experiments and characterized expression of synaptic markers. L.T. carried out live imaging of circuit activity in wild type and mutant mice. S.C, S.B., L.T., J.T.T., K.S., and S.L.Z analyzed the data. S.C., S.B., K.S. and S.L.Z. wrote the paper with input from L.T. and J.T.T.

## METHODS

### Mice

Mouse breeding and husbandry procedures were carried out in accordance with UCLA’s animal care and use committee protocol number 2009-031-31A, at University of California, Los Angeles. Mice were given food and water *ad libitum* and lived in a 12-hr day/night cycle with up to four adult animals per cage. Only virgin male C57BL/6J wild-type mice were used for single nuclei sequencing and FISH experiments in this study.

For genetic analysis of *Igsf9b*, mice used in immunohistochemistry and 2-photon imaging experiments were naive subjects with no prior history of participation in research studies. All live imaging was performed on mice expressing jGCaMP7f in V1B neurons. GCaMP expression was induced by AAV pGP-AAV-syn-jGCaMP7f-WPRE intracortically injected into Igsf9b WT and KO mice. WT (*Igsf9b^+/+^*) and KO (*Igsf9b^-/-^*) mice were bred from *Igsf9b^+/-^* mice graciously gifted by the Krueger-Burg lab. These mice were originally obtained by their lab from Lexicon Pharmaceuticals (Thee Woodlands, TX, U.S.A.; Omnibank clon 281214, generated through insertion of the Omnibank gene trap vector 48 into Igsf9b gene in Sv129 ES cells). The commercial version of this mouse has since sold to Taconic Biosciences (1 Discovery Drivee, Suite 304, Rensselaer, NY 12144) (https://www.taconic.com/knockout-mouse/igsf9b-trapped). The mice were backcrossed onto C57BL/6J background for at least 6 generations by the Krueger-Burg lab and confirmed to be null KO in Babaev et al 2018 (https://doi.org/10.1038/s41467-018-07762-1). Genotyping was performed on P6-P9 pups, and genotypes of pups were identified by PCR that was outsourced to Transnetyx (transnetyx.com). Plots for NR and DR mice in Figure 7G, J and K were from unpublished results in Tan et al., 2020. A total of 14 mice, both male (9) and female (5) were used in 2-photon imaging. P21 WT: 3 males and 1 female. P21 KO: 1 male and 2 females. P36 WT: 4 males and 1 female (1 female overlaps with P21 WT). P36 KO: 1 male and 4 females (2 females overlap with P21 KO).

### Visual deprivation experiments

Mice that were dark-reared were done so in a box covered from inside and outside with black rubberized fabric (ThorLabs Cat# BK5) for 7-17 days (P21-P28 or P21-P38) or 9 days (P8-P17) before being euthanized. The dark box was only opened with red light on in the room (mice cannot perceive red light). Mice that were dark-light reared were first dark reared for 7 days from P21 to P28 in the dark, and then transferred back to the mouse room to receive 8 hours of ambient light prior to euthanasia.

### V1 dissection to obtain single nuclei

Normally-reared mice were dissected at P8, P14, P17, P21, P28, and P38. Isoflurane was used for anesthetization and mice were euthanized by cervical dislocation. Dark-reared mice were dissected at P28 and P38. Dark-light reared mice were dissected at P28 after exposure to 8 hr ambient light. For each age or condition group, 30 mice were dissected: 15 for each biological replicate of single-nucleus(sn) RNA-sequencing. Mice were anesthetized in an isoflurane chamber, decapitated, and the brain was immediately removed and submerged in Hibernate A (BrainBits Cat# HACA). While the dissection was aimed to target V1b, the region enriched for binocular neurons, due to the small size of this region, the dissection invariably captured neighboring V1 tissue. Therefore, we refer to the tissue as V1. Extracted brains were placed on a metal mold and the slice containing V1 was isolated by inserting one blade 0.5 mm posterior to the lambdoid suture and a second blade 1.5 mm further anterior (2 spaces on the mold). This slice was removed and lowered to Hibernate A in a 60cc petri dish, which was placed on a ruler under a dissecting microscope. The midline was aligned with the ruler and the first cut was bilaterally 3.3 mm out from the midline. The second cut was 0.7 mm medial to the first cut. The cortex was peeled off the underlying white matter. The V1 piece with a total of 1 mm cortex depth by 1.5 mm thickness was transferred to a dish containing 600 µl of RNAlater (Thermo Fisher Cat# AM7020) and kept on ice until dissections were complete. Dissected tissues were then kept in RNAlater at 4°C overnight and transferred to -20°C the next day. Tissue was stored this way for up to 1 month prior to being processed for snRNA-seq.

### Droplet-based single-nucleus(sn) RNA-sequencing

For each biological replicate, dissected V1 regions from 15 mice were removed from RNAlater, weighed, then chopped with a small blade on a cleaned slide on top of a cooling cube. Tissue was then transferred to a dounce homogenizer chilled to 4°C and denounced slowly 30 times with a tight pestle in 1 ml of homogenization buffer containing 250mM Sucrose, 150mM KCl, 30mM MgCl_2_, 60mM Tris pH 8, 1 µM DTT, 0.5x protease inhibitor (Sigma-Aldrich Cat# 11697498001), 0.2 U/µl RNase inhibitor, and 0.1%TritonX. All solutions were made with RNase-free H_2_O. Each sample was filtered through a 40 μm cell strainer and then centrifuged at 1000g for 10 minutes at 4°C. The pellet was resuspended in the homogenization buffer and an equal volume of 50% iodixanol was added to the resuspended pellet to create 25% iodixanol and nuclei mix. This mix was layered upon 29% iodixanol and spun at 13,500g for 20 minutes at 4°C. The supernatant was removed and the pellet was washed in a buffer containing 0.2 U/µl RNAse inhibitor, PBS (137 mM NaCl, 2.7 mM KCl, 8 mM Na_2_HPO_4_, and 2 mM KH_2_PO_4,_ pH 7.4), 1% bovine serum albumin, and then filtered over a 40 μm filter and centrifuged at 500g for 10 minutes at 4°C. The pellet was resuspended and filtered with two more 40 μm filters, cells counted on a hemocytometer and then diluted to 700-1200 nuclei/mm^3^. Nuclei were re-counted on a 10X automated cell counter. Nuclei were further diluted to the optimal concentration to target capturing 8000 cells per channel.

Nuclei from each biological replicate were split into two and run separately on two channels of 10X v3, targeting 8,000 cells per channel. We refer to these as library replicates. For each experiment, we performed two biological replicates towards a total of four library replicates. The two biological replicates were processed on different days. Sequencing was performed using the Illumina NovaSeq™ 6000 Sequencing System (S2) to a depth of ∼30,000 reads per cell. All library preparation and sequencing were performed at the UCLA’s Technology Center for Genomics & Bioinformatics (TCBG) core.

### Single-molecule fluorescent *in situ* hybridization (smFISH)

C57/BL6J mice were anesthetized in isoflurane at ages ranging from P8 to P38 and then perfused transcardially with heparinized PBS followed by 4% paraformaldehyde (PFA) diluted in PBS and adjusted to pH 7.4. Following perfusion, the brains were collected and postfixed for 24h at 4°C in 4% PFA, and then cryoprotected sequentially in 10%, 20%, and 30% sucrose in PBS solution until the brain sank. Brains were then frozen in OCT using a methylbutane and dry ice bath and stored at -80°C until time of sectioning. Brains were cut into 15 μm thick coronal sections at -22/-20°C using a cryostat (Leica CM 1950) and single sections were collected in a charged microscope slide in ascending order from the frontal to the occipital region starting in V1. For localization of the visual cortex V1 and binocular zone of V1, coordinates from ^61^ were used. Sections were stored at -80°C until further processing. For all FISH experiments, coronal sections were selected to be from a similar anatomical region within V1 when comparing conditions or ages.

Multiplex FISH was performed following ACD Biology’s Multiplex RNAscope v2 assay (Advanced Cell Diagnostics, cat# 323110). Briefly, thawed sections were baked at 60°C, post-fixed for 1 hr at 4°C in 4% PFA, and then dehydrated in sequential ethanol treatments followed by H_2_O_2_ permeabilization and target retrieval. Protease III treatment was used, then application of probes and sequential amplification and fluorophore development fluorophores (Akoya Biosciences cat# FP1487001KT, FP1488001K, FP1497001KT). Slides were counterstained with 1 ug/ml 4,6-diamidino-2-pheenylindole (DAPI, Sigma cat #D9542) and mounted with Prolong Gold (Thermo Fisher Scientific cat# P36930). RNAscope probes used include: *Igsf9b* (cat# 832171-C3), *Mdga1* (cat#546411, 546411-C2), *Nlgn2* (cat# 406681). Cdh13 (cat # 443251-C3), *Chrm2* (cat # 495311-C2), *Deptor* (cat #481561 - C3), *Gad1* (cat3 400951-C2), *Slc17a7* (cat# 416631-C2, 416631, 416631-C3), *Trpc6* (cat# 442951), *Tshz2* (cat# 431061-C1). Each time point or condition had three to four biological replicates comprising brain sections from different mice. NR mice at P8, P14, P17, P21, P28, and P38, DR mice at P17, P28, and P38, and DL mice at P28 were used.

### Immunolabeling for synaptic markers

Immunolabeling for VGLUT1 and GAD65 on P5 brains was performed on perfusion-fixed brains that underwent the same preparation as for smFISH. Brains were sectioned to 15 μm sections. Sections were then incubated for 24 hr with anti-VGLUT1 (guinea pig polyclonal Millipore Sigma Cat# AB5905) and anti-GAD65 (mouse monoclonal Millipore Sigma Cat#MAB3521R) diluted 1:500 in blocking solution (10% NGS in 0.3% PBST), washed 3x times in PBS, and then incubated for 2 hr with goat anti-mouse 488 (Invitrogen Cat# A11029) and goat anti-guinea pig 568 (Invitrogen Cat#A11075) both diluted 1:500 in blocking solution.

Immunolabeling for synaptic markers in IGSF9B KO vs WT experiments were performed on perfusion-fixed brains sectioned to 40 µm and preserved in aliquots of antifreeze (42.8g Sucrose Fisher Cat # S25590B, 0.33g of MgCl2.6H2O Sigma Cat#M2670, 312.5g (250 mL) glycerol Sigma Cat#G7757, 25mL 10X PBS, total to 500 mL w/ ddH2O). On the day of the experiment, tissues were washed 4 times at 15 minutes per wash from antifreeze using 0.3% PBST, blocked with 10% goat serum in PBST, incubated with primary antibody at 4C for 2 nights (∼44 hours). Samples were washed 4 times, 15 minutes each in 1X PBS, then secondary antibody diluted in blocking solution was added for 2 hr at room temperature. Samples were washed 4 times at 10minutes per wash in 1X PBS then stained with 1:10k DAPI for 15 minutes, washed for 10 min in PBS, and then mounted with Prolong Gold. Primary antibodies used include: anti-VGLUT1 (guinea pig polyclonal Millipore Sigma Cat# AB5905), anti-VGLUT2 (Guinea pig polyclonal Synaptic Systems Cat#135404, anti-GAD65 (mouse monoclonal Millipore Sigma Cat#MAB3521R), anti-NLGN2 (guinea pig polyclonal Synaptic Systems Cat#129205), anti-GABRA1 (rabbit polyclonal Synaptic Systems Cat#224203), anti-GABRG2 (guinea pig polyclonal Synaptic Systems Cat#224004 (anti-GPHN (mouse monoclonal Synaptic Systems Cat#147011), anti-VGAT (rabbit polyclonal Synaptic Systems Cat#131002), anti-PSD95 (rabbit polyclonal Invitrogen Cat#VH307495), anti-SSCAM (rabbit polyclonal Sigma Aldrich Cat#2441). Secondary antibodies used include goat anti-guinea pig 566 (Invitrogen Cat#A11075), goat anti-guinea pig 647 (Life Technologies Cat#A21450), goat anti-mouse 488 (Invitrogen Cat#A11029), goat anti-guinea pig 568 (Invitrogen Cat#A11075), goat anti-rabbit 488 (Life Technologies Cat#A11008), goat anti-mouse 568 (Invitrogen Cat#A11031), goat anti-rabbit 647 (Invitrogen Cat#A21244), goat anti-rabbit 568 (Invitrogen Cat#A11011).

### Confocal imaging

Images were acquired on a Zeiss 880 confocal microscope at 20X and 40X magnification. Each image was 1024 pixels x 1024 pixels in size. For 20X and 40X images, this corresponded to a 0.4 μm x 0.4 μm and 0.2 μm x 0.2 μm coverage per frame, respectively. Vertically tiled 20X images were acquired covering the entire cortex, as well as 40X horizontal tiled images to cover L2/3 only. Z-stacks covered the entire 15 μm section. *Mdga1* and *Ccbe1,* both L2/3 -markers, were used as markers to assess the cortical depth covered by each 40X image. For each 40X frame starting at layer 2, one frame covered the depth of L2/3 based on *Mdga1* and *Ccbe1* signals. For immunolabeling experiments, images were taken using a confocal microscope with 63X magnification, imaged on both sides of the brain in L2/3 of V1 based on anatomical markers. One z-stack comprising 5 optical sections spanned the entire 15 μm section imaged. For thick-section (40 µm) immunohistochemical sections, 15 µm z-stacks were taken at 63X using two vertically tiled frames that correspond to a 0.13 by 0.26 µm imaged zone of L2/3.

### Imaging quantification

3D z-stacked images were z-projected on FIJI version 2.1.0/1.53c. The entire z-stack covering the slide was projected into a 2D image with maximum intensity. 20X images were tiled using DAPI and *Slc17a7* channels (when available) as guides through linear blending to capture the entire cortical thickness. 40X and 63X images were processed as is. Maximum-projected images were entered into CellProfiler using a custom pipeline modified from the original SABER-FISH pipeline ^62^. Modifications were made to detect up to four imaging channels ^63^. CellProfiler was used to perform nuclear and cell segmentation, as well as puncta counting. Nuclear segmentation was done by using DAPI and cellular segmentation was done by taking a fixed radius of 5 pixels around the nucleus. For downstream computation, nuclear segmentation results were used. Segmented images had nuclear boundaries as well as individual puncta married in an overlay color with original image items in gray. All segmented images were inspected to ensure no aberration in segmentation or puncta calling.

After segmentation and puncta calling, data were analyzed in R using custom scripts to compare nuclear mRNA counts (i.e., number of puncta) between time points and conditions. For cell type experiments, cells were sorted into types based on mRNA counts of marker genes. Briefly, cells were ranked based on their mRNA counts of each gene and visualized as a scatter plot of counts vs. rank. The knee of this plot was located ^64^. The mRNA count value at the knee was chosen as the cutoff for cell type assignment. Quantification of protein puncta in immunolabeling experiments also used Cell Profiler by adapting the same pipeline developed to count mRNA puncta. Protein puncta were quantified per image and normalized by the number of nuclei segmented in that image (Figure S7B-F). In Figure S7G, the unique peri-nuclear somatic distribution of NLGN2 enabled its quantification using a 20-pixel boundary around the nucleus and counting the puncta that fell in that boundary. This allowed for quantifying NLGN2 protein puncta per cell.

### Surgery and AAV injection

All epifluorescence and two-photon imaging experiments were performed through chronically implanted cranial windows. Mice aged P10-11 (for P21 imaging) or P25-26 (for P36 imaging) were administered with carprofen analgesia prior to surgery, anesthetized with isoflurane (5% for induction; 0.75–1.5% during surgery), mounted on a stereotaxic surgical stage via ear bars and a mouth bar. Their body temperature was maintained at 37°C via a heating pad. The scalp and connective tissue were removed, and the exposed skull was allowed to dry. Then a thin layer of Vetbond was used to cover the exposed skull and wound margins. Once dry, a thin layer of dental acrylic was applied around the wound margins, avoiding the region of skull overlying V1. A metal head bar was affixed with dental acrylic caudally to V1. A 3mm circular piece of skull overlying binocular V1 on the left hemisphere was removed after using high-speed dental drill to thin the bone along the circumference of this circle. Care was taken to ensure that the dura was not damaged at any time during drilling or removal of the skull.

Local AAV injection into binocular V1 took place after the skull was removed. Exposed brain was submerged in normal saline during injection. AAV was diluted in 1xPBS that contains 2.5% mannitol (w/v) to a final titer of 6.7∼7.5×10^12^ genomes per ml. Mannitol was used to increase the viral spread (Mastakov et al., 2001). For both age groups, virus was injected at least 10 days before imaging. Virus injection was done using a glass micropipette (tip diameter: 19-25 μm) and Nanoject III (Drummond Scientific Company) attached on Scientifica PatchStar Micromanipulator (Scientifica) controlled with LinLab2 (Scientifica). Injection site was at 3 mm lateral from the midline and 1 mm rostral from lambda. Injections occurred at three depths: 470, 340 and 210 µm below the pial surface. At each depth, 65 cycles of injection were done, with each cycle injecting 5 nL at 5 nL/s speed, with 10 second intervals between cycles. Thus, 325 nL of AAV was injected at each depth, and 975 nL was injected into V1B in total. After virus injection, a sterile 2.5 mm diameter cover glass was placed directly on the exposed dura and sealed to the surrounding skull with Vetbond. The remainder of the exposed skull and the margins of the cover glass were sealed with dental acrylic. Mice were then recovered on a heating pad. When alert, they were placed back in their home cage. Carprofen was administered daily for 3 days post-surgery. Mice were left to recover for at least 10 days prior to imaging. Mice injected at P10-11 would also be imaged at P36 if their cranial windows remained clear.

### Mapping of binocular area of the primary visual cortex

The precise location of the binocular region in V1 on the left hemisphere for each mouse was identified using low magnification, epifluorescence imaging of jGCaMP7f signals. For all mice, visual areas were mapped the day before imaging. Briefly, jGCaMP7f was excited using a 470nm light-emitting diode. A 27-inch LCD monitor (ASUS, refreshed at 60 Hz) was positioned such that the binocular visual field fell in the center of the monitor. The screen size was 112 deg in azimuth and 63 deg in elevation and the monitor was placed 20 cm from the eyes. A contrast reversing checkerboard (checker size 10×10 degree) bar windowed by a 1D Gaussian were presented along the horizontal or vertical axis to both eyes (Figure S7H). The checkerboard bar drifted normal to its orientation and swept the full screen width in 10 sec. Both directions of motion were used to obtain an absolute phase map along the two axes. Eight cycles were recorded for each of the four cardinal directions. Images were acquired at 10 frames per second with a PCO edge 4.2 sCMOS camera using a 35mm fixed focal length lens (Edmund optics, 35mm/F1.65, #85362, 3mm field of view). The camera focused on the pial surface. The visual areas were obtained from retinotopic maps of azimuth and elevation. The binocular area of the primary cortex was defined as the region of primary visual cortex adjacent to the higher visual area LM (Figure S7I).

### Two-photon calcium imaging

Two-photon imaging was targeted to the binocular area of V1 using a resonant/galvo scanning two-photon microscope (Neurolabware, Los Angeles, CA) controlled by Scanbox image acquisition software (Los Angeles, CA). A Coherent Discovery TPC laser (Santa Clara, CA) running at 920 nm focused through a 16x water-immersion objective lens (Nikon, 0.8 numerical aperture) was used to excite jGCaMP7f. The objective was set at an angle of 10-11 degrees from the plumb line to reduce the slope of the imaging planes. Image sequences (512×796 pixels, 490×630 μm) were captured at 15.5 Hz at a depth of 120 to 320μm below the pial surface on alert, head-fixed mice that were free to run on a 3D-printed running wheel (14cm diameter). A rotary encoder was used to record the rotations of this running wheel. Three planes that were well separated in depth and covered the top, middle and bottom of L2/3 were imaged per mouse (Figure S7J). To measure responses of neurons to each eye separately, an opaque patch was placed immediately in front of one eye when recording neuronal responses to visual stimuli presented to the other eye.

### Visual stimulation during 2-photon imaging

On the same screen that was used for visual area mapping, a set of sinusoidal gratings with 18 orientations (equal intervals of 10 degrees from 0 to 170 degrees), 12 spatial frequencies (equal steps on a logarithmic scale from 0.0079 to 0.1549 cycles per degree) and 8 spatial phases were generated in real-time by a Processing sketch using OpenGL shaders (see https://processing.org). These static gratings were presented at 4 Hz in full screen in pseudo-random sequence with 100% contrast (Figure 7A). Imaging sessions were 15 min long (3600 stimuli in total), thus each combination of orientation and spatial frequency appeared 16 or 17 times. Transistor-transistor logic signals were used to synchronize visual stimulation and imaging data. The stimulus computer generated these signals, and these were sampled by the microscope electronics and time-stamped by the acquisition computer to indicate the frame and line number being scanned at the time of the TTL.

### Analysis of two-photon imaging data

#### Image processing

Movies for either eye from the same plane were processed together using a standard pipeline consisting of movie concatenation, motion correction, cell segmentation and ROI signal extraction using Suite2p (https://suite2p.readthedocs.io/). ROIs determined for each experiment were inspected and confirmed visually (Figure S7K). Neuronal spiking was estimated via non-negative temporal deconvolution of the extracted ROI fluorescence signal using Vanilla algorithm (Figure S7L) (Berens et al., 2018). Subsequently, fluorescent signals and estimated spiking for each cell were split into separate files corresponding to the individual imaging session for each eye. Each imaging plane was processed independently.

#### Calculation of response properties

##### Identification of visually responsive neurons using SNR

Signal to noise ratio (SNR) was used to identify neurons with significant visual responses (tuned neurons). SNR for each neuron was calculated based on the optimal delay of the neuron. Optimal delay was defined as the imaging frame after stimulus onset at which the neuron’s inferred spiking reached maximum. To calculate SNR, signal was the mean of standard deviations of spiking to all visual stimuli around the optimal delay (4-6 frames, thus ∼0.323 sec, after stimulus onset; see Figure7H), and noise was this value at frames well before or after stimulus onset (frames –2 to 0, and 13 to 17). Neurons whose optimal delays occurred outside of the time-locked stimulus response window of 3 to 7 frames after stimulus onset (padded by ±1 frame around the 4-6 frame range used above) were spontaneously active but visually unresponsive. They were untuned neurons and had SNR values close to 1 (Figure 7I). The SNR values of these untuned neurons were normally distributed (mean = 1.0±0.03) over a narrow range. Untuned neurons with optimal delays naturally occurring in the 3-7 frame time window can be distinguished from visually responsive neurons by SNR. This SNR threshold was defined at 3 standard deviations above the mean SNR of the above-mentioned normal distribution (See the vertical dashed lines in Figure 7K). SNR values were calculated separately for responses to the ipsilateral or contralateral eye. A neuron is monocular if its SNR for one eye, but not the other, was above the threshold (Figure 7C). A neuron is binocular if its SNR for either eye was above the threshold (Figure 7D). A neuron is untuned if its SNR for neither eye was above the threshold.

##### Tuning kernel for orientation and spatial frequency

The estimation of the tuning kernel was performed by fitting a linear model between the response and the stimulus. Cross-correlation maps were used to show each neuron’s inferred spiking level to each visual stimulus (orientation and spatial frequency) and were computed by averaging responses over spatial phases. The final tuning kernel of a neuron was defined as the correlation map at the optimal delay (Figure 7B).

##### Orientation preference

We used vertical slices of the tuning kernel through the peak response and calculated the center of mass of this distribution as orientation preference. Orientation preference calculation:

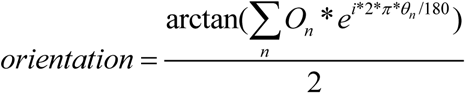

O_n_ is a 1×18 array, in which a level of estimated spiking (O_1_ to O_18_) occurs at orientations θ_n_ (0 to 170 degrees, spaced every 10 degrees). Orientation is calculated in radians and then converted to degrees.

##### ΔOrientation for binocular neurons

For a binocular neuron, *Ori_contra_* is the neuron’s orientation preference to contralateral eye and *Ori_ipsi_* is the orientation preference to ipsilateral eye. *ΔOrientation* was calculated as

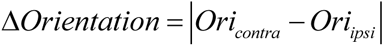

If the value of *ΔOrientation* is above 90 (e.g., |170-10|=160), then the actual value for the difference of orientation preferences to two eyes is 180- *ΔOrientation* (180-160=20).

#### Quantification and Statistical Analysis for 2-photon imaging

A power analysis was not performed a-priori to determine sample size. All statistical analyses were performed in MATLAB (https://www.mathworks.com/), using non-parametric tests with significance levels set at α < 0.05. Bonferroni corrections for multiple comparisons were applied when necessary. Mann-Whitney U-tests (Wilcoxon rank sum test) or two-sample Kolmogorov-Smirnov tests were used to test differences between two independent populations.

### Computational analysis of single-nucleus transcriptomics data

#### Alignment and quantification of gene expression

Fastq files with raw reads were processed using Cell Ranger v3.1.0 (10X Genomics) with default parameters. The reference genome and transcriptome used was GRCm38.92 based on Ensembl 92, which was converted to a pre-mRNA reference package by following Cell Ranger guidelines. Each single-nucleus library was processed using the same settings to yield a gene expression matrix (GEM) of mRNA counts across genes (columns) and single nuclei (rows). Each row ID was tagged with the sample name for later batch correction and meta-analysis. We henceforth refer to each nuclear transcriptome as a “cell.”

#### Initial pre-processing of normally reared samples to define classes, subclasses, and types

This section outlines the initial transcriptomic analysis of data from normally reared samples. Unless otherwise noted, all analyses were performed in Python using the SCANPY package ^58^. The complete computational workflow is illustrated in Figure S1D.

1. Raw GEMs from 23 snRNA-seq libraries were combined: 6 ages, 2 biological replicates per age and 2 library replicates per biological replicate except for P38, where one of the technical replicates failed quality metrics at the earliest stage of processing. This resulted in a GEM containing 184,936 cells and 53,801 genes.
2. We then generated scatter plots of the number of transcript molecules in each cell (n_counts), the percent of transcripts derived from mitochondrially encoded genes (percent_mito), and the number of expressed genes (n_genes) to identify outlier cells. Cells that satisfied the following conditions were retained: 700 < n_genes < 6500, percent_mito < 1%, and n_counts < 40,000. Only genes detected in more than 8 cells were retained for further analysis. This resulted in a GEM of 167,384 cells and 30,869 genes.
3. Cells were normalized for library size differences. Transcript counts in each cell were rescaled to sum to 10,000 followed by log-transformation. For clustering and visualization, we followed steps described previously ^65^. Briefly, we identified highly variable genes (HVGs), z-scored expression values for each gene, and computed a reduced dimensional representation of the data using principal component analysis (PCA). The top 40 principal components (PCs) were used to compute a nearest-neighbor graph on the cells. The graph was then clustered using the Leiden algorithm ^66^ and embedded in 2D via the Uniform Manifold Approximation and Projection (UMAP) algorithm ^67^.

#### Additional filtering and class assignment

The analysis above yielded 42 clusters (Figures S1E-F). Canonical marker genes for cortical classes and subclasses were used to annotate these clusters (Figure S1G**, Table S1**). We then used Scrublet ^28^ to identify doublets (Figure S1H). Clusters that expressed markers of two or more classes or contained more than 50% doublets were labeled “Ambiguous” (Figure S1I). Removal of ambiguous clusters and doublets in the dataset resulted in a GEM containing 147,236 cells by 30,868 genes.

For further analysis, this matrix was subsetted by cell class (glutamatergic neurons, GABAergic neurons, and non-neuronal cells) and age (P8, P14, P17, P21, P28 and P38) into 18 separate GEMs (Figure S1D).

#### Identification of cell types within each class by age

Each of the 18 GEMs were separately clustered using the procedure described above with one modification. Following PCA, we used Harmony ^60^ to perform batch correction. The nearest-neighbor graph was computed using the top 40 batch-corrected PCs.

Each GEM was then iteratively clustered. We began by clustering cells using the Leiden algorithm, with the resolution parameter fixed at its default value of 1. As before, UMAP was used to visualize the clusters in 2D. Through manual inspection, small clusters with poor quality metrics or ambiguous expression signatures were discarded, likely representing trace contaminants that escaped detection in the earlier steps. The remaining clusters were annotated by subclass based on canonical expression markers (**Table S1,** Figure 1D). Next, we performed a differential expression (DE) analysis between each cluster and other clusters in its subclass. If a cluster did not display unique expression of one or more genes, it was merged with the nearest neighboring cluster in the UMAP embedding as a step to mitigate over-clustering. This DE and merging process was repeated until each cluster had at least one unique molecular signature (Figures S3A-C). We refer to the final set of clusters as types.

#### Workflow for supervised classification analyses

To assess transcriptomic correspondence of clusters across ages or between rearing conditions, we used XGBoost, a gradient boosted decision tree-based classification algorithm ^59^. In a typical workflow, we trained an XGBoost (version 1.3.3) classifier to learn subclass or type labels within a “reference” dataset, and used it to classify cells from another, “test” dataset. The correspondences between cluster IDs and classifier-assigned labels for the test dataset are used to map subclasses or types between datasets. The classification workflow is described in general terms below and applied to various scenarios in subsequent sections.

Let *R* denote the reference dataset containing *N_R_* cells grouped into *r* clusters. Let *T* denote the test dataset containing *N_T_* cells grouped into *t* clusters. Here, each cell is a normalized and log-transformed gene expression vector ***u*** ∈ *R* or ***v*** ∈ *T*. The length of ***u*** or ***v*** equals the number of genes. Based on clustering results, each cell in *R* or *T* is assigned a single cluster label, denoted cluster(***u***) or cluster(***v***). cluster(***u***) may be a type or subclass identity, depending on context.

The main steps are as follows:

1. We trained multi-class XGBoost classifiers *C_R_^0^* and *C_R_^T^* on *R* and *T* independently using all 30,868 genes as features. In each case, the dataset was split into training and validation subsets. For training we randomly sampled 70% of the cells in each cluster, up to a maximum of 700 cells per cluster. The remaining “held-out” cells were used for evaluating classifier performance. Clusters with fewer than 100 cells in the training set were upsampled via bootstrapping to 100 cells in order to improve classifier accuracy for the smaller clusters. Classifiers achieved a 99% accuracy or higher on the validation set. XGBoost parameters were fixed at the following values:

1. ‘Objective’: ‘multi:softprob’
2. ‘eval_metric’: ‘mlogloss’
3. ‘Num_class’: *r* (or *t*)
4. ‘eta’: 0.2
5. ‘Max_depth’: 6
6. ‘Subsample’: 0.6
2. When applied to a test vector ***c***, the classifier *C_R_^0^* or *C_R_^T^* returns a vector *p = (p_1_, p_2_, …)* of length *r* or *t*, respectively. Here, *p_i_* represents the probability value of predicted cluster membership within *R* or *T,* respectively. These values are used to compute the “softmax” assignment of ***c***, such that cluster(***c***) = *arg max_i_ p_i_* if *arg max_i_ p_i_* is greater than *1.2*(1/r)* or 1.2*\*(1/t)*. Otherwise ***c*** is classified as ‘Unassigned’.
3. Post training, we identified the set of top 500 genes based on average information gain for each *C_R_^0^* and *C_R_^T^*. These gene sets are denoted *G_R_* and *G_T_*.
4. Using the common genes *G = G_R_ ⋂ G_T_*, we trained another classifier *C_R_* on 70% of the cells in R, following the procedure outlined in 1. As before, the performance of *C_R_* was evaluated on the remaining 30% of the data.
5. Finally, we trained a classifier *C_R_* on 100% of the cells in R. *C_R_* was then applied to each cell ***v*** ∈ *T* to generate predicted labels cluster(***v***).

#### Comparing transcriptomic signatures of developmental V1 to adult V1/ALM subclasses (Tasic et al., 2018)

We used the aforementioned classification workflow to evaluate the correspondence between V1 subclasses in this work (Figure 1D) and those reported in a recent study of the adult V1 and motor cortex (ALM) ^20^. We trained a classifier on the V1/ALM subclasses and used it to assign an adult label to each V1 cell collected in this study. A confusion matrix was used to visualize the correspondence between developmental V1 subclasses and V1/ALM subclasses at adulthood (Figure S1J). This correspondence served as a proxy to evaluate the overall conservation of subclass-specific transcriptomic signatures across developmental stages (developing vs. adult), RNA source (single-nucleus vs. single-cell), platform (3’ droplet-based vs. full-length plate-based), and region (V1 vs. V1/ALM).

#### Inferring temporal association between V1 types using supervised classification

##### Relating types across time

The supervised classification workflow was used to relate cell types identified at each pair of consecutive ages within each class (5 x 3 = 15 independent analyses). In each case, the classifier was trained on the older age dataset and applied to each cell in the younger age dataset. Thus, each cortical cell at the younger age possessed two type labels, one identified via clustering of cells at that age and the other based on a classier trained at the next age. Assessing the correspondence between these labels enabled us to link cell types between consecutive ages (e.g., P8-P14, P14-P17 and so on) and track their maturation across development

##### Quantification and visualization of cluster correspondence

The correspondences between types throughout development were visualized using Sankey flow diagrams (Figures 2E, S2F-G). In the case of glutamatergic neurons, for example, inspecting the Sankey flow diagrams revealed that L2/3 and 4 types mapped more diffusely across time than L5 and 6 types, suggesting subclass specific differences in maturation. We quantified such subclass-specific differences using three methods,

1. We computed the adjusted rand index (ARI) between the cluster labels and classifier-assigned labels. The ARI ranges from 0 and 1, with extremes corresponding to random association and perfect (i.e., 1:1) mapping, respectively. Negative values are possible for the ARI but were not observed in our data. The ARI was computed using the function `sklearn.metrics.adjusted_rand_score().` ARI values were computed for each pair of consecutive ages (e.g., P8 and P14) within each subclass (e.g., L2/3). ARI differences between glutamatergic subclasses were visualized as bar plots (Figure 2F). The analysis was repeated for GABAergic and non-neuronal cells (Figure S2H-I).
2. We computed for each type the F1 score, which is a measure of a classifier’s effectiveness at associating cells within a type to their correct type label. Its value ranges from 0 to 1, with extremes corresponding to no association and perfect association between transcriptome and type label, respectively. The F1 score was computed for each type at each time point using the function `sklearn.metrics.f1_score()`. Values were grouped by subclass to visualize differences (Figures S2J-L). This analysis showed that in addition to exhibiting poor temporal correspondence, L2/3 and L4 types were also less transcriptomically distinct than L5 and L6 types at any given time point (Figure S2J). Subclasses within GABAergic and non-neuronal cells did not exhibit such striking differences (Figures S2K-L).
3. We assessed the sensitivity of each subclass’ clustering results to the clustering resolution parameter of the Leiden algorithm, which controls the number of output clusters. The clustering resolution was increased from 1 to 2. We computed the ARI between the clusters identified at each value of the resolution parameter and the baseline clusters computed at a resolution value of 1. The ARI was computed for the clusters within each subclass at each time point separately. L2/3 and L4 clustering was more sensitive to changes in the resolution parameter than the clustering in L5 and L6 (Figure 2G).

#### Analysis of visual deprivation experiments

##### Separation of major cell classes

In visual deprivation experiments, snRNA-seq profiles were collected from cortical samples of mice dark-reared from P21-P28 (P28DR), dark-reared from P21-P38, (P38DR) and dark-reared from P21-P28 followed by 8 hours of ambient light stimulation. Overall, 12 GEMs from these three experiments were combined and preprocessed (4 libraries per experiment) using the steps described above for normally reared samples. The numbers of cells prior to pre-processing were 43,234, 36,373 and 31,815 for P28DR, P38DR and P28DL respectively. The final numbers of high-quality cells reported were 24,817, 25,671, and 26,575, respectively.

##### Comparing DR and DL clusters to NR types using supervised classification

To examine cell type correspondence between visual deprivation and normally reared experiments, we used supervised classification as described above. Classifiers were trained on P28NR and P38NR types, and cells from P28DL, P28DR, and P38DR were mapped to the corresponding NR age. The resulting confusion matrices were visualized as dot plots, and the ARI was computed for types within each subclass (Figure S5).

#### Differential gene expression analysis

Differential expression (DE) was performed in multiple settings to identify genes enriched in specific classes, subclasses, types, or rearing conditions. We used the `scanpy.tl.rank_genes_groups()` function and Wilcoxon rank-sum test in the scanpy package for statistical comparisons ^58^. While searching for genes enriched in a particular group of interest, only those expressed in >20% of cells in the tested group were considered.

The results of the DE analyses were used in the following contexts: (1) To assess the quality of cell populations identified in the initial analysis, where each cluster in Figure S1F was compared to the rest. Clusters that did not express a unique signature or those that express markers known to be mutually exclusive were removed; (2) To identify novel subclass markers (Figure 2B, Figure S2D-E). This was accomplished by comparing each subclass against the rest; (3) To identify type-specific markers within each subclass (Figure S3A-C). Here, each type was compared to other types of the same subclass; and (4) To identify gene expression changes as a result of visual deprivation. We performed DE between NR and DR (both ways) subclasses (Figure 6A, Figure S6A).

#### Identification of genes showing graded expression among L2/3 types

We compared each L2/3 type to the other two (e.g., A vs B and C) to identify 287 type-specific genes at fold change > 2 and p-value < 10^-10^ (Wilcoxon test). The expression levels of these genes were z-scored, and we used *k*-means clustering to identify *k*=7 groups based on their pattern of expression among the three types (Figure S6A). The optimal number of groups was identified using the elbow method. Five of the seven groups, containing 217 genes, showed graded expression differences that could be classified into one of the following patterns based on visual inspection: A > B > C (77 genes), A < B > C (36 genes), C > A > B (9 genes), C > B > A (85 genes) and A > C > B (10 genes). The remaining X genes were expressed in a digital fashion that fell into one of two groups: C > B = A (35 genes) and A > B = C (35 genes). Thus, approximately 75% of the DE genes among L2/3 types are expressed in a graded fashion.

#### Pseudo-spatial inference of gene expression in L2/3

FISH experiments targeting the three L2/3 glutamatergic type markers revealed that type A resides at the top (near the pia), type B in the middle, and type C at the bottom of L2/3, bordering L4 (Figure 3). Surprisingly, this relative positioning of A, B, and C types was mirrored in the UMAP embedding. We therefore hypothesized that the UMAP coordinates of a neuron may serve as a proxy for the approximate relative position of its soma in the tissue and used this to calculate the expected spatial expression profiles of genes in each dataset.

In a given scenario, we marked the “A” and “C” cells furthest from each other on the UMAP space as the “root” and the “leaf” and assumed that these represent the top and bottom of L2/3 respectively. We used diffusion pseudo-time (DPT) ^68,69^ to order all L2/3 cells relative to the root cell. DPT and similar methods have been used previously to order cells based on their developmental state (i.e., pseudo-time); we have used it in this context to infer “pseudo-spatial” position based on the observed correspondence described above. Pseudo-spatial positions for cells were close to 0 at the top, where type A begins, and gradually increased through types B and C, reaching the maximum normalized value of 1 at the end of L2/3 in UMAP space. We performed this pseudo-spatial analysis for L2/3 neurons in each of the six normally reared samples.

For the DR and DL datasets, where the spatial organization and transcriptomic profiles are disrupted, a root cell was randomly selected from the beginning of L2/3 in UMAP space (e.g., a cell from the edge of cluster “L2/3_1” was chosen for P28DR) (Figure 4B). Finally, to visualize the expression of gradient genes as a function of pseudo-spatial position (Figure 6F, H), we averaged the expressions along bins of pseudo-spatial location that contained as many cells as ∼10% of a given dataset.

#### Identification of temporally regulated genes

This analysis was repeated separately for each of L2/3, L4, L5, and L6. Of the 30,868 genes in the data, we considered only those expressed in more than 20% of the cells in at least one of the six time points. This resulted in 6339, 5746, 6096, and 5428 genes for further analysis in L2/3, L4, L5, and L6, respectively. We first computed the average expression strength of every gene at each of the six time points. Here, the average expression strength *E_g,t_* of gene *g* at age *t* is defined as follows,

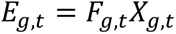

where *F_g,t_* is the fraction of cells at age *t* that express gene *g* and *X_g,t_* is the mean transcript counts of *g* among cells with non-zero expression. We only considered genes that satisfied the following condition,

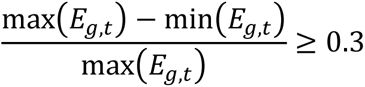

resulting in 2594, 2410, 2190, and 2192 genes for further analysis in L2/3, L4, L5, and L6, respectively. Next, to identify genes that showed significant temporal variation, we z-scored each *E_g_* vector and randomly shuffled the temporal identities of the cells. We then recomputed a randomized analog of *E_g,t_*, which we call 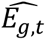. We then defined for each gene *g* a deviation score between the actual and randomized expression vectors,

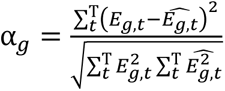

Here, *T* = 6 is the number of time points and the denominator acts as a normalizing factor; we observed a bias towards highly expressed genes in its absence. High values of *α_g_* indicate that the observed temporal pattern of expression is significantly different from the randomized pattern. We picked 855 genes for further analysis that had *α_g_* > 0.2. This threshold was chosen by computing an empirical null distribution for *α_g_* using two randomizations 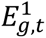 and 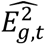. The 99.9^th^ percentile value of *P_null_*(*α_g_*) was 0.05, making *α_g_* = 0.2 a conservative threshold. Finally, we counted the number of temporally differentially expressed (tDE) genes identified in each layer (Figure S3H)

#### Separation of cell classes and subclasses using Seurat

In addition to clustering each time separately in SCANPY, Seurat (version 3.1, ^32^ was used to cluster data from all times and conditions together. This analysis was done to evaluate class and subclass level clustering, and to provide a framework to broadly check gene expression for FISH experiments in all subclasses at all times collectively. Seurat clustering was performed using two methods with similar final results. In the log-normalization based method, data were log normalized and scaled to 10,000 transcripts per cell, with 2000 variable genes used. In the generalized linear model method “SCTransform” ^70^, normalization was used with 3000 variable genes. In both methods cells with fewer than 1000 or over 6000 genes or >1% mitochondrial content were filtered out. PCA was performed and unsupervised clustering was applied to the top 80 PCs. Major cell type markers from ^1^ and ^20^ were used to assign class and subclass designations to clusters. Clusters having two or more major markers were discarded as “doublet/debris” clusters, and clusters that were solely composed of one or two replicates were also discarded as debris clusters. In both log-normalization and SCT clustering by Seurat, the P8 cortico-cortical projecting excitatory neurons clustered separately from similar subclass neurons of later time points. Thus, P8 was clustered separately, and cell IDs from P8-only clustering were used to re-label the corresponding P8 cells in the full dataset. Class and subclass level clustering results matched SCANPY-based results (Figure S2J).

#### Differential gene expression analysis using Seurat

The Seurat-based clustering results were primarily used to assess subclass-level differentially expressed genes. Gene signatures of each cell subclass at different time points were identified with the `FindMarkers` function, performing pairwise time or condition comparisons and by comparing one time point to the average of others (a second method only used normally reared datasets). Genes were considered if they were present in 10% of cells, 0.25 log fold enriched (1.28 fold-change or more), and had a Benjamini-Hochberg corrected *P*<0.05. Of these, genes that were 0.4 log fold enriched (1.5-fold change or more) were classified as enriched.

### Data and software availability

Computational scripts detailing snRNA-seq analysis reported in this paper along with the raw expression matrices and processed h5ad files are available at https://github.com/shekharlab/mouseVC. All raw and processed snRNA-seq datasets reported in this study will be made publicly available via NCBI’s Gene Expression Omnibus (GEO). All data and custom software for imaging analysis will be made available upon request.

## SUPPLEMENTAL FIGURES, TITLES, AND LEGENDS

**Figure S1.**
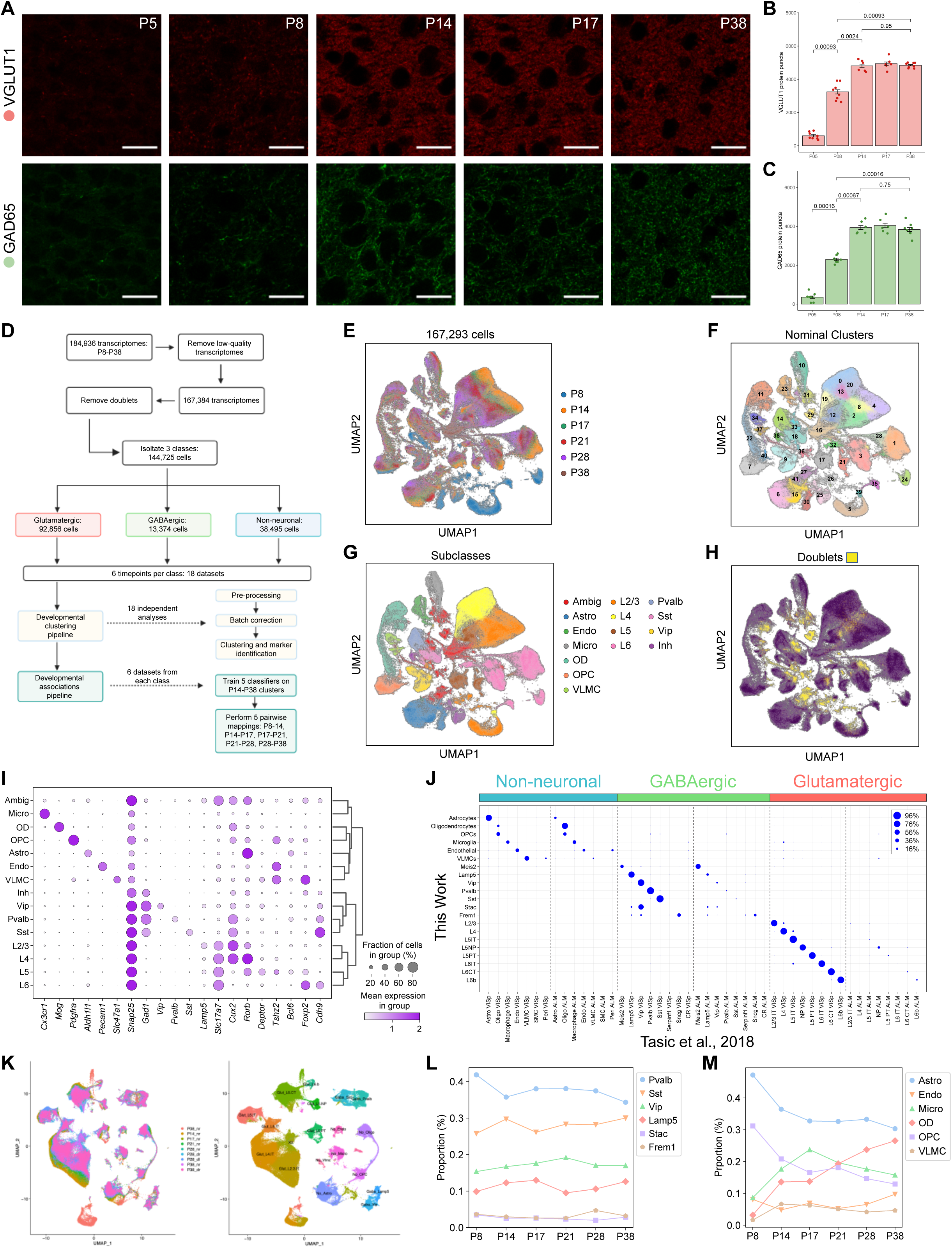
Expression of GABAergic and glutamatergic markers with age, snRNA-seq data pre-processing, and comparison to transcriptomic signatures of adult visual and motor cortices (Tasic et al., 2018), related to Figure 1. A. Immunohistochemical analysis of the expression of VGLUT1 (a glutamatergic marker) and GAD65 (a GABAergic marker) with age. Panels show single confocal images of VGLUT1 (red) and GAD65 (green) during postnatal development in L2/3 of V1 in wild-type mice. Ages are indicated on the top right corner panels in the top row. Scale bar, 20 μm. B. VGLUT1 protein puncta quantification over time, n = 3-4 mice per age, quantified by slide imaged. Horizontal bars show p-values corresponding to a comparison of the number of puncta between each pair of ages using the Wilcoxon rank-sum test. Bar heights denote mean value, error bars are ± SEM. C. Same as B, for GAD65 protein puncta. D. A graphical summary of the computational pipeline used to define cell classes, subclasses, and types at each age in normally reared (NR) mice and to analyze the maturation of types. A similar pipeline was used for analyzing the data obtained from dark-rearing (DR) and dark-light adaptation (DL) experiments described in Figure 4. See **Methods** for details. E. Uniform Manifold Approximation (UMAP) visualization of V1 transcriptomes prior to removing doublets. Individual nuclei are colored by age. F. Same as E, colored by transcriptomically defined clusters. This nominal clustering was used to identify and discard doublets and contaminants (**Methods**). G. Same as E, with nuclei colored by their subclass identity. Clusters in F were assigned to subclasses based on known gene markers (**Table S1).** H. Same as E, with computationally identified doublets highlighted in yellow. The doublets were identified using Scrublet, a nearest-neighbor classification framework that uses the data to simulate multiplets ^28^. A post hoc analysis verified that the computationally identified doublets co-expressed markers from distinct classes and subclasses (**Table S1)** and were discarded prior to further analysis. I. Dot plot showing marker genes (columns) that distinguish subclasses (rows) identified in panel G. The group labeled “Inh” is an admixture of inhibitory neuronal subclasses defined by *Lamp5, Stac,* and *Frem1* that were not separated at this stage. The size of each dot represents the fraction of cells in each subclass with non-zero expression and its color denotes the average normalized expression level. J. Confusion matrix showing transcriptomic correspondence between subclasses identified in this study (rows) and subclasses (columns) reported in a scRNA-seq survey of the adult primary visual cortex (V1) and the anterior lateral motor cortex (ALM) ^20^. The size of each dot represents the proportion of each row mapped to an adult cortex subclass based on an XGBoost classifier trained on the adult cortex. Each row of the matrix is normalized to sum to 100%. The specific pattern of mapping indicates that the transcriptomic correspondence at the level of classes and subclasses was robust despite differences in RNA source (nuclei vs. cells), age (P8-P38 vs. P70+), and sequencing methods (e.g., droplet-based and plate-based). Glutamatergic and GABAergic subclasses map specifically to their V1 counterparts, reflecting regional specificity. Non-neuronal cells, on the other hand, mapped promiscuously, suggesting shared transcriptional programs between V1 and ALM for this class. K. UMAP visualization of subclass separation when the same dataset is analyzed using Seurat (**Methods**). (*Left*) colored by sample ID, (*right*) colored by subclass. X_0_ denotes low quality cells that were discarded. L. Line plots showing that the relative proportions of GABAergic neuronal subclasses remain stable with age despite significant variation in the number of nuclei collected (**Table S2**). M. Line plots showing relative proportions vs. age for non-neuronal subclasses.

**Figure S2.**
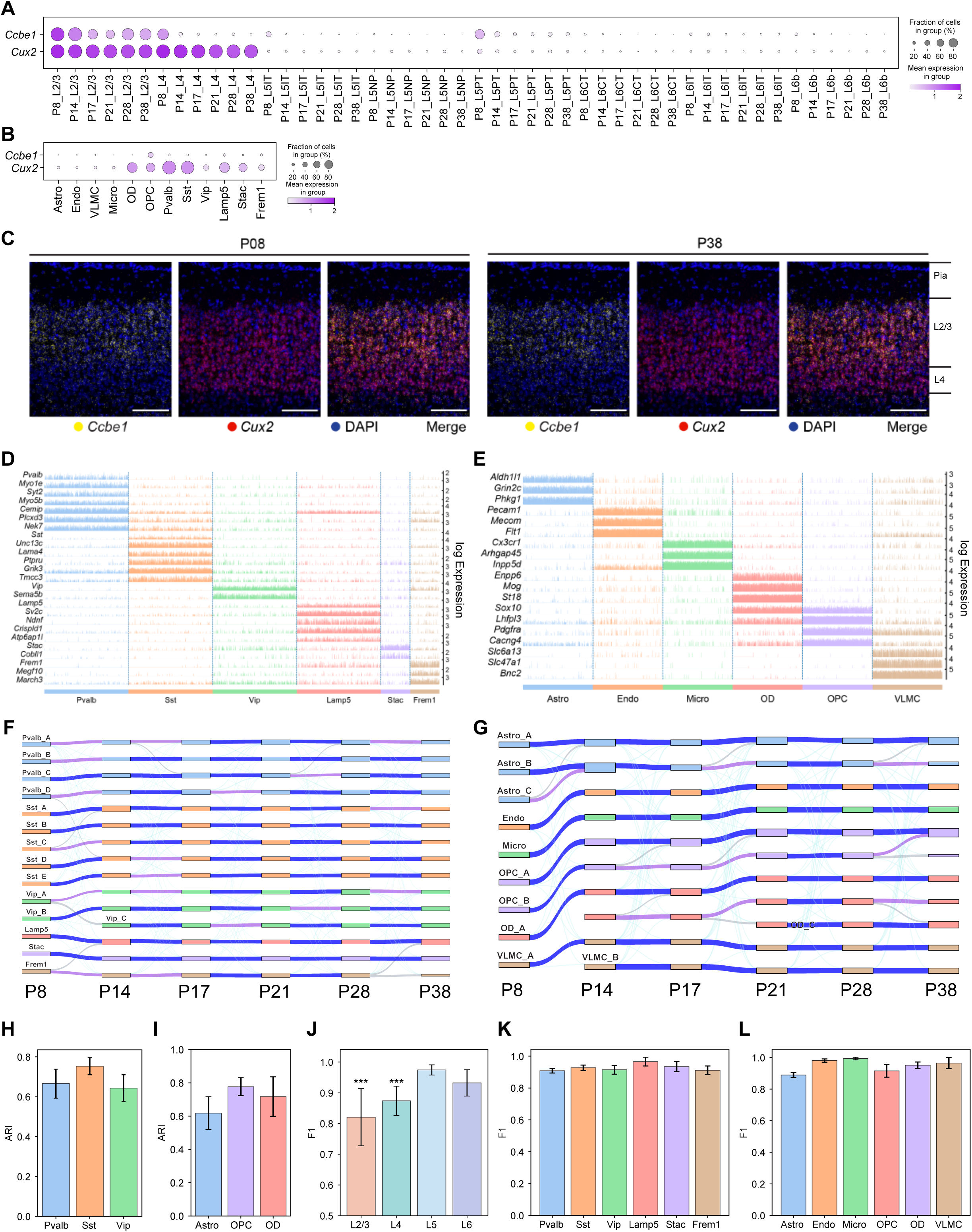
Additional data related to transcriptomic maturation of V1 cell types, related to Figure 2. A. *Ccbe1* is specific at all ages to L2/3 glutamatergic neurons (note transient expression in L5PT at P8), whereas *Cux2* is expressed in both L2/3 and L4 glutamatergic neurons throughout development. B. *Cux2* is also expressed in GABAergic neurons and some non-neuronal subclasses, while *Ccbe1* is not. Thus, *Ccbe1* is a bona fide L2/3 glutamatergic neuron-specific marker. C. FISH images from coronal sections show that *Cux2* is expressed in L2/3 and L4 while *Ccbe1* is selectively expressed in L2/3 glutamatergic neurons at P8 and P38. Scale bar, 100 μm. D. Tracks plot showing subclass-specific markers (rows) in inhibitory neurons (columns). For each gene, the scale on the y-axis (right) corresponds to normalized, log-transformed transcript counts detected in each cell. Columns are grouped and colored based by subclass (annotation bar, bottom). In addition to the well-known subclasses marked by *Pvalb*, *Sst* and *Vip,* we also identify subclasses of GABAergic neurons marked by the selective expression of *Lamp5*, *Stac* and *Frem1*. These subclasses were collapsed together in the group labeled “Inh” in Figure S1I. E. Same as panel D for non-neuronal subclasses. F. Sankey graph showing the transcriptomic maturation of V1 GABAergic types. Representation as in Figure 2E. G. Same as F for non-neuronal types. H. Adjusted Rand Index (ARI) values quantifying temporal specificity during maturation of inhibitory types within each subclass. ARI is a measure of similarity between two ways of grouping the data, with values ranging from 0 (no correspondence) to 1 (perfect correspondence). For each subclass at each age, ARI values are computed by comparing the cluster identity of cells with the identity assigned by a multiclass classifier trained on the subsequent age. Individual bars denote the three inhibitory subclasses containing more than 2 types each. Subclasses *Lamp5*, *Stac* and *Frem1* are not shown as these could not be satisfactorily subdivided into constituent types in this study. Bar heights, mean ARI computed across pairs of consecutive ages; error bars, standard deviation. I. Same as panel H for non-neuronal subclasses. Subclasses Micro and Endo are not included as they each contain only one type. J. F1 score values quantifying the degree of transcriptomic separation among types within each glutamatergic layer. Each bar represents the average F1 score calculated for types from each layer across all ages. *P*\*\*\*<0.0001 for comparison between L2/3 and L5/L6 and L4 and L5/6. K. Same as panel J for GABAergic types. L. Same as panel J for non-neuronal types.

**Figure S3.**
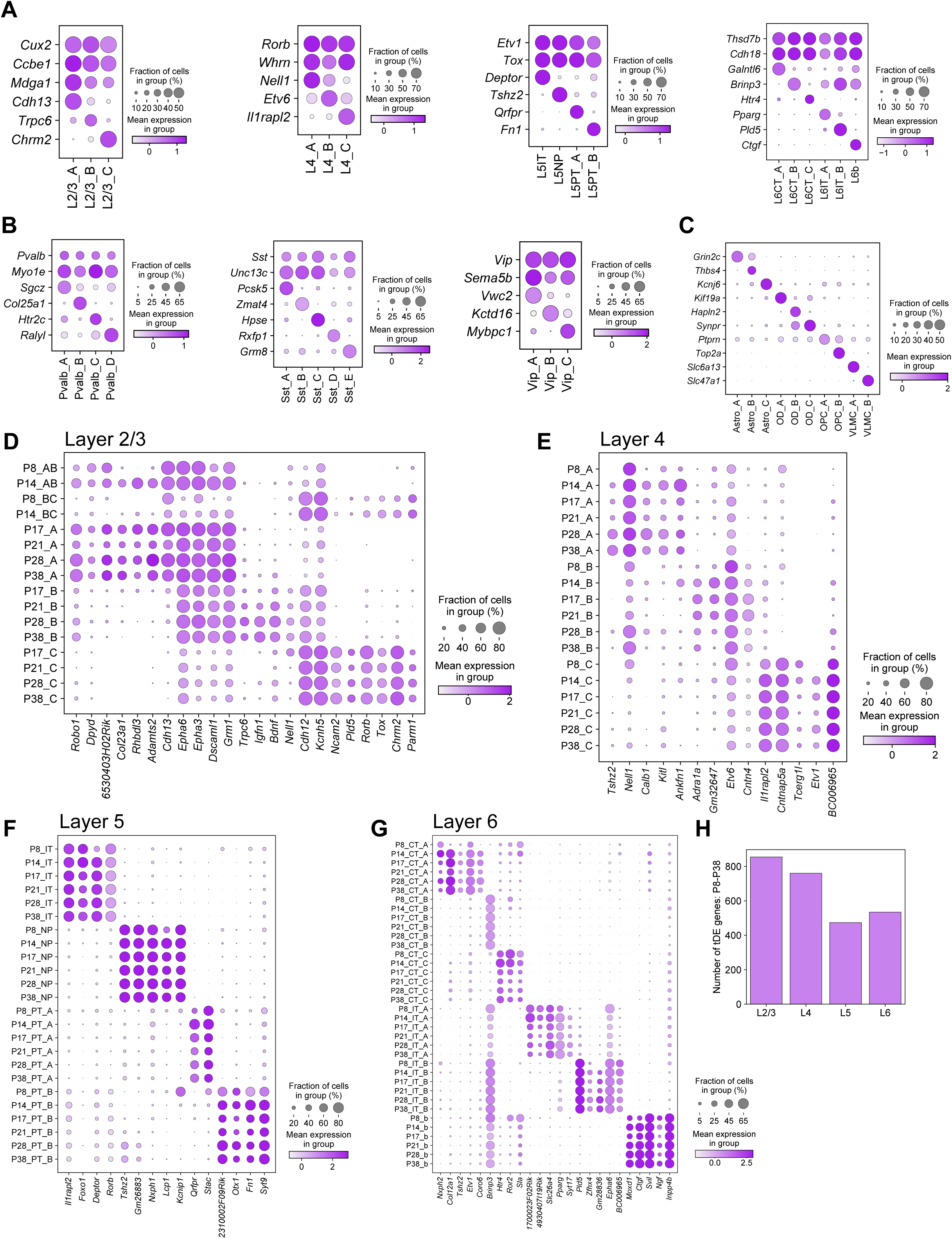
Cell type-specific markers for glutamatergic, GABAergic, and non-neuronal subclasses, related to Figure 2. A. Dot plot showing markers (rows) for glutamatergic types (columns) within each subclass (panels, left to right). Within each subclass panel, the top two genes (rows) represent markers shared by all types, while the remainder are type-specific markers. Expression levels are computed by pooling cells from all ages. Representation as in Figure S1I. B. Same as A, for GABAergic types within subclasses (panels, left to right). C. Same as A, for non-neuronal types. D. Dot plot showing that type-specific markers for L2/3 glutamatergic neurons vary with age. Also shown are markers for the precursor types AB and BC at P8 and P14, which are expressed in types A, B and C at later ages (P17-P38). E. Dot plot showing that type-specific markers for L4 glutamatergic neurons also vary with age. F. Dot plot showing that type-specific markers for L5 glutamatergic neurons are stable with age G. Dot plot showing that type-specific markers for L6 glutamatergic neurons are stable with age. H. Bar plot showing the number of temporally differentially expressed (tDE) genes found in each layer. The elevated number of tDE genes in L2/3 and 4 is consistent with their late specification (see Figure 2). The selection criteria for tDE genes were identical for all layers (see **Methods**).

**Figure S4.**
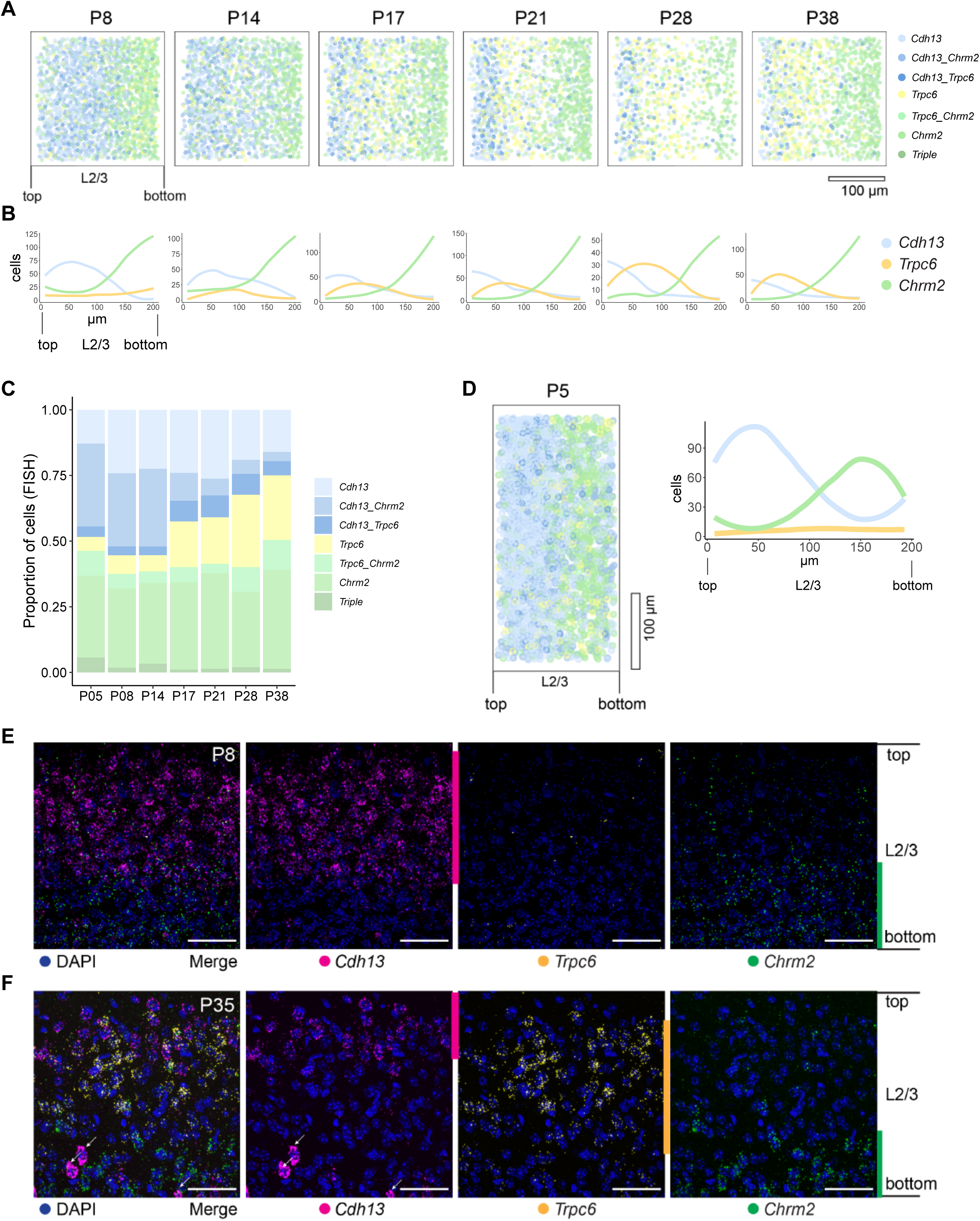
*In situ* analysis of L2/3 glutamatergic neuron types in normally reared mice, related to Figure 3. A. Pseudo-colored representation of imaged cells (**Methods**) within L2/3 expressing one or more of the type markers *Cdh13, Trpc6,* and *Chrm2* across the six ages. Each panel represents overlaid FISH images of 18 V1 regions from three mice imaged at 40X with the pial-ventricular axis oriented horizontally from left to right. Each imaging frame is 208 μm x 208 μm and is imaged starting at the top of L2/3 to the end of the imaging frame covering about the usual thickness of L2/3 in one frame. As pia could not be imaged within the same frame at 40X, alignment of overlaid images is less optimal compared to 20X images in the main figure. During quantification, this leads to an underrepresentation of *Cdh13^+^* cells at the top of the imaging frame in 40X compared to 20X images. B. Line tracings quantifying number of cells per bin at each position along the pial to ventricular axis (x-axis) divided into 14 equally spaced bins, corresponding to panel A. 0 on the x axis is the region of L2/3 closest to pia. C. Relative proportions of cells within each expression group defined in panel A as quantified by FISH. Total number of cells analyzed: P5, 2106; P8, 3734; P14, 2937; P17, 3102; P21, 2078; P28, 2078; and P38, 2775. Scale bar, 100 µm. D. Same as panels A and B, at P5. E. 40X FISH images of L2/3 at P8 labeled with the three type markers *Cdh13*, *Trpc6*, and *Chrm2* that become evident after P17. *Cdh13* and *Chrm2* are selectively expressed between the precursor types AB and BC at earlier ages. *Trpc6* is not expressed at this early stage. Pial to ventricular axis is oriented vertically from top to bottom. Scale bar, 50 µm. F. 40X FISH images of L2/3 at P35 labeled with the three type markers *Cdh13*, *Trpc6*, and *Chrm2* showing sublayering along the pial to ventricular axis (top to bottom). Arrows denote inhibitory neurons expressing *Cdh13*. Scale bar, 50 µm.

**Figure S5.**
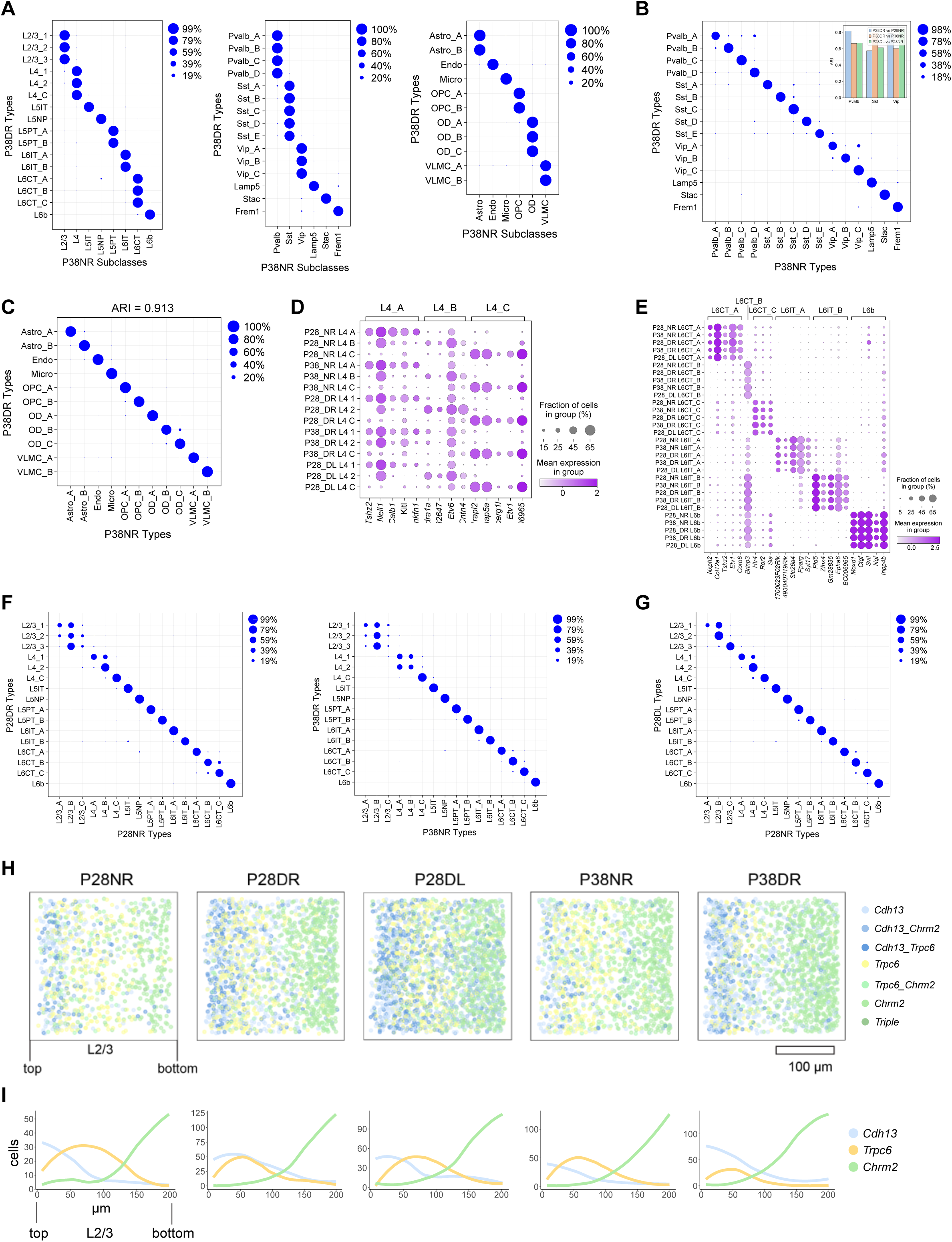
Transcriptomic changes in V1 cell type diversity in dark-rearing experiments, related to Figure 4. A. Confusion matrices from supervised classification analyses showing that transcriptomically-defined clusters in dark-reared (DR) mice (rows) map to the correct subclass in normally reared (NR) mice (columns). This shows that the transcriptome-wide signatures that define the major subclasses are not disrupted by visual deprivation. Panels correspond to glutamatergic (*left*), GABAergic (*middle*) and non-neuronal (*right*) subclasses at P38. Results are similar when comparing P28DR vs. P28NR or P28DL vs. P28NR (data not shown). B. Confusion matrix from a supervised classification analysis showing that GABAergic clusters in P38DR mice (rows) transcriptomically correspond 1:1 to GABAergic types in P38NR mice (columns). Results are similar when comparing P28DR vs. P28NR and P28DL vs. P28NR (data not shown). Together these results show that the transcriptomic signatures that define types within GABAergic neurons are not disrupted by visual deprivation. Inset, bar plot of the adjusted rand index (ARI) showing high transcriptomic correspondence between P28DR, P38DR and P28DL types and NR types at the same age. C. Same as panel B, showing the 1:1 transcriptomic correspondence of non-neuronal types between P38DR vs P38NR. ARI value is indicated on top. Results are similar for P28DR vs. P28NR and P28DL vs. P28NR (data not shown). Thus, the transcriptomic signatures that define non-neuronal types are not disrupted by visual deprivation. D. Dot plot showing expression patterns of L4 type markers (columns) in NR types at P28 and P38, and DR and DL clusters at the same ages (rows). Markers are grouped by those specific for NR types L4_A, L4_B and L4_C (annotation bar, top). E. Same as panel D for L6 types. L6 types in DR and DL experiments show a 1:1 correspondence with the NR types (panels D and E) and are therefore named accordingly. A similar 1:1 mapping is not possible for L2/3 and L4 clusters in DR and DL experiments because of the disruption of type-specific gene expression signatures. F. Confusion matrices showing the transcriptomic correspondence between glutamatergic clusters in DR vs. types in NR at P28 (left) and P38 (right). With the exception of the three types in L2/3 and two types in L4, all the remaining types have a 1:1 match with a DR cluster. Thus, visual deprivation selectively impacts type-specification within L2/3 and L4 among glutamatergic types. G. Confusion matrix showing the transcriptomic correspondence between glutamatergic clusters in DL vs. types in NR at P28. While the patterns are similar to panel D, L2/3 and L4 clusters show an increased correspondence, reflecting partial recovery of cell type specific signatures. H. Pseudo-colored representation of imaged cells (**Methods**) within L2/3 expressing one or more of the type markers *Cdh13, Trpc6,* and *Chrm2* across five combinations of age and conditions, indicated on top. Each panel represents overlaid FISH images of 15-18 V1 regions from three mice imaged at 40X, with the pial-ventricular axis oriented horizontally from left to right. Each imaging frame is 208 μm x 208 μm and is imaged starting at the top of L2/3 to the end of the imaging frame covering about the usual thickness of L2/3 in one frame. I. Line tracings quantifying number of cells per bin at each position along the pial to ventricular axis (x-axis), divided into 14 equally spaced bins, corresponding to panel H. 0 on the x-axis is the region of L2/3 closest to pia. Total number of cells analyzed in H, I: P28, 933; P28DR 1671; P28DL, 2148; P38NR, 1419; and P38DR, 1784.

**Figure S6.**
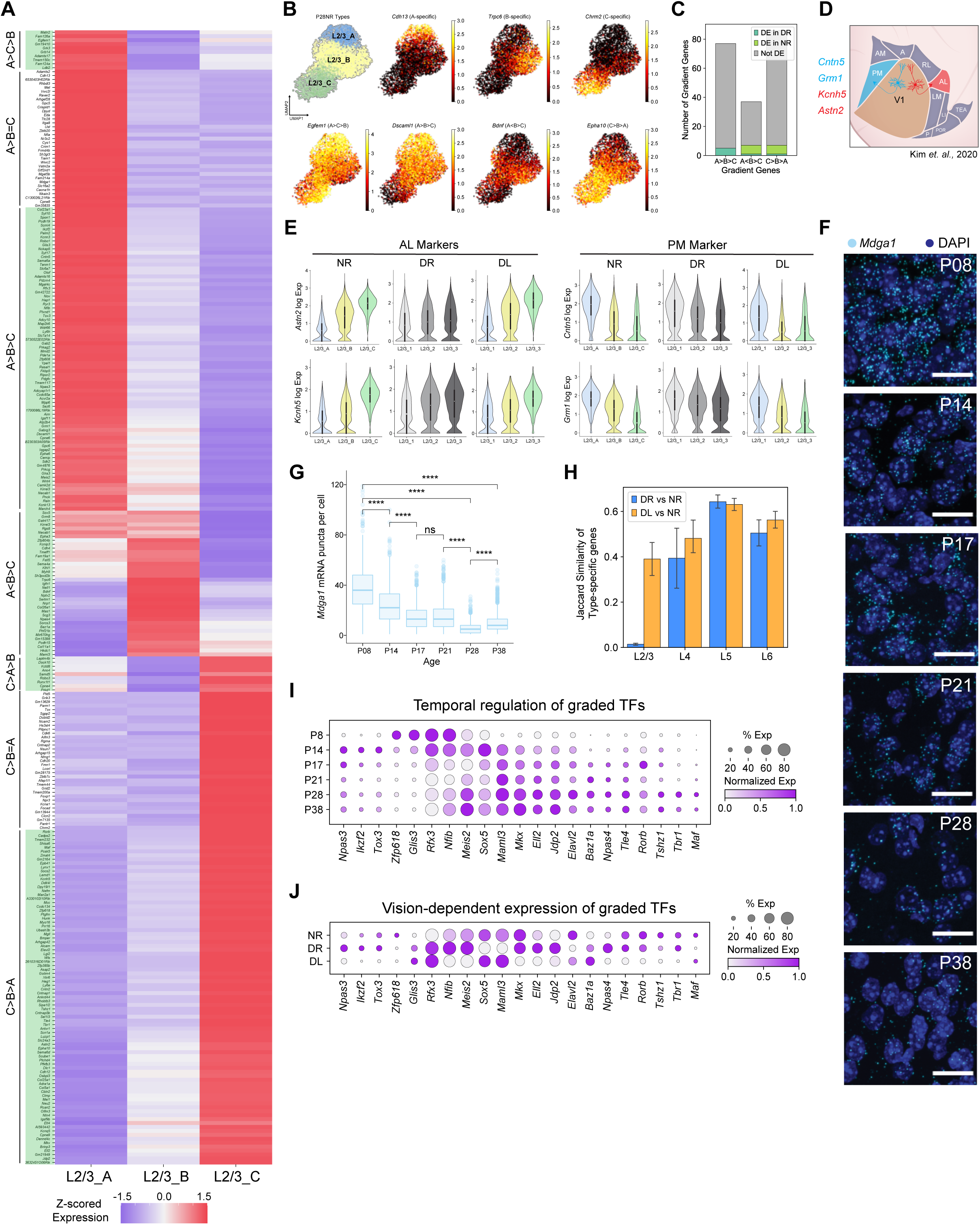
Graded gene expression among L2/3 types, and selective maturation of L2/3 types, related to Figure 6. A. Type-specific genes for L2/3 types A, B and C (columns) are predominantly expressed in a graded (i.e., not a digital) fashion among the types. Each type was compared to the other two, and only genes with a fold-change (FC) cutoff >2 were selected (FDR< 0.05 by Wilcoxon rank-sum test). Colors denote z-scored expression levels across the three types. Genes (rows) are grouped by expression patterns among the types (e.g., A> B> C). The analysis was performed at P28NR, and the pattern of expression is similar at P38NR (data not shown). Green shading highlights graded genes, which are shown in Figure 6A. B. UMAP feature plots of L2/3 neurons at P28NR with cells colored based on their expression of selected DE genes from panel A. The leftmost panel on the top row shows the locations of the three types L2/3_A, L2/3_B, and L2/3_C. The remaining panels on the top row show three genes that are digitally expressed among the three L2/3 types. The bottom panels show four genes expressed in a graded fashion among the three L2/3 types. C. Bar plot showing that only a small fraction of graded (shaded) genes from A are differentially expressed when L2/3 cells are compared in bulk between DR and NR mice at P28 (fold-change>2, *P-value* < 10^-10^ by Wilcoxon rank-sum test). Thus, visual deprivation does not change the average expression levels of these genes but disrupts their graded patterning. D. Schematic highlighting the anterolateral (AL) and posteromedial (PM) higher visual areas. Genes preferentially expressed by L2/3 neurons projecting to AL or PM are indicated. Colors identified by retrograde labeling experiments are listed ^46^. E. Violin plots showing expression of markers enriched in AL (*left*) and PM (*right*) projecting neurons from Kim et al., 2020 in L2/3 cell types in NR mice vs. L2/3 cell clusters in DR and DL mice at P28. The graded expression in NR types is disrupted in DR clusters and partially recovered in DL clusters. F. FISH images showing the expression of *Mdga1* mRNA over time in V1. Three animals per time point, six images per animal. Scale bar, 20 μm. G. Box plot quantifying expression in panel F. Wilcoxon Rank Sum Test, **** p <0.0001. Number of cells quantified: P8,1191; P14,1011; P17, 1389; P21, 1729; P28, 1277; and P38, 1588. H. Bar plot showing the dependence of visual experience (DR and DL) on cell type specific genes within each layer. Each bar represents the Jaccard similarity of type-specific genes between different experience conditions. This was computed for each type within a layer separately and the mean and standard deviation are shown. While the gene expression signatures within all four layers are different for mice with different visual experience conditions, the effect is most dramatic for L2/3 (see L2/3 DR vs NR). In addition, DL significantly recovers the similarity of L2/3 signatures to those found in NR. I. Dot plot showing the temporal regulation of transcription factors (TFs) found in Figure 6A. Expression levels shown for L2/3 neurons. J. Same illustration as panel I across the conditions P28NR, P28DR, and P28DL.

**Figure S7.**
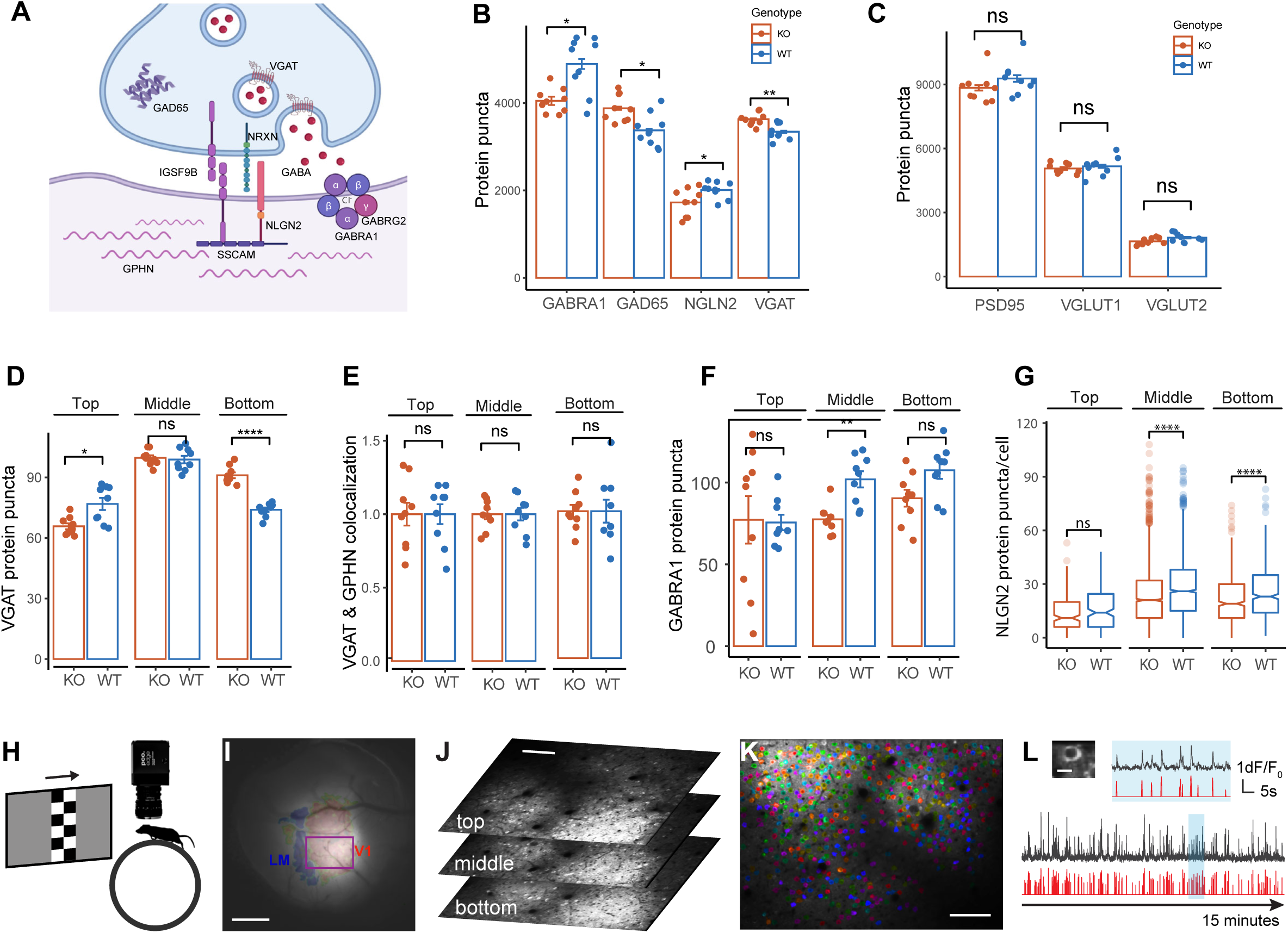
Immunohistological characterization of synaptic markers in *Igsf9b* KO, mapping V1b, and measuring receptive field tuning via 2-photon calcium imaging of awake mice, related to Figure 7. A. Schematic of molecules at an inhibitory synapse. GAD65, VGAT, and NRXN are presynaptic. IGSF9B is both pre- and post- synaptic and binds homophilically. The intracellular component of Igsf9b interacts with SSCAM in the postsynaptic compartment, which also interacts with the intracellular domain of NLGN2 and, thereby, may stabilize NRXN-NLGN interaction at the cell surface. GABA receptors are post-synaptic. They comprise 5 subunits including subunits GABRA1 and GABRG2. B. Levels of GABRA1, GAD65, NLGN2, and VGAT protein in L2/3 of WT (n = 9) vs KO (n = 9) mice at P35-38. Y-axis shows total puncta counted within a 130 x 260 μm zone of L2/3 within V1 extending 260 µm from the top of L2/3 to the base of L2/3, and across by 130 μm. *p < 0.05. **p <0.005 by Wilcoxon Rank Sum test. Exact P values: GABRA1, p = 0.0078; GAD65, p = 0.0056 VGAT, p = 0 .0028; NLGN2, p = 0.05. C. No significant difference in expression of excitatory synapse markers in KO vs WT, 9 mice per group, measured in L2/3 of V1 as in Figure 7B. D. VGAT protein as a function of L2/3 depth in KO (n = 9) vs WT (n = 9) mice. Top, p = 0.0078; Middle, p = 0.55; and bottom, p = 4.1×10^-5^. Top, middle, and bottom regions of layer 2/3 are defined based on FISH results for type A, B, C markers. Top corresponds to the first 25 μm of L2/3 and bottom corresponds to the lower 50 μm with the middle being everything in between. E. VGAT and GPHN colocalization as a function of L2/3 depth in KO (n = 9) vs WT (n = 9) mice. F. GABRA1 as a function of L2/3 depth in KO (n = 9) vs WT (n = 9) mice, normalized by number of nuclei in the corresponding zone. Top region ns, Middle p = 0.00078, Bottom p = 0.063 (ns but trending). G. NLGN2 protein puncta per cell, in WT (n = 9) vs KO (n = 9) mice at P35-39. Total quantified cells: 2945 from 9 mice in KO and 2786 cells from 9 mice in WT. Due to the perisomatic localization of Nlgn2, its level of expression was quantified on a cell by-cell basis by using a fixed 20-pixel ring around the nucleus as the cell border. **** p < 0.00005 by Wilcoxon Rank Sum test. Exact P values: Top ns. Middle p = 1.06 x 10^-23^, Bottom p = 3.96x 10^-5^. H. Mapping visual cortical areas to localize the binocular region of V1 (V1b) using low-magnification epifluorescence imaging of jGCaMP7f evoked responses to checkerboard bars that were both drifting and flashing. I. Example image of a cranial window highlighting V1b and its border with higher-order lateromedial (LM) area. The purple rectangle delineates the field of view used for 2-photon imaging. Scale bar, 0.5 mm. J. Three planes of in vivo 2-photon calcium imaging of neurons within layer 2/3 in the field of view in panel I. The three imaging planes cover the top, middle and bottom sub-laminae in layer 2/3. Scale bar, 100 μm. K. Overlay of motion-corrected average fluorescence image and segmentation of neurons for the middle plane in J. Scale bar, 100 μm. L. Top left: a neuron expressing jGCaM7f in the bottom plane of panel J. Tuning kernel of this neuron was shown in Figure 7B. Scale bar, 10 μm. Bottom: the raw (black) and temporally deconvolved (red) jGCaMP7f signal from this neuron for 15 minutes of visual stimulation to the contralateral eye. The region in blue is expanded above horizontally to show more details of the signal.

## SUPPLEMENTAL TABLES

**Table S1. List of canonical markers used to identify neuronal and non-neuronal subclasses.**

**Table S2. Number of cells present in each of the 18 NR and 9 DR/DL datasets.**

